# Immunohistochemistry-compatible gel-assisted mass spectrometry imaging

**DOI:** 10.64898/2026.07.31.742113

**Authors:** Maddison C. Hibbard, Yat Ho Chan, Helen P. Dobbelmann, Koralege C. Pathmasiri, Stephanie M. Cologna, Ruixuan Gao

## Abstract

While lipids are fundamental to cellular signaling and structure, probing native lipid compositions and distributions across complex tissue architectures at single-cell resolution remains challenging due to inherent trade-offs in spatial resolution, sensitivity, and molecular coverage in current spatial lipidomics methods. Here, we report a multimodal imaging approach that seamlessly integrates gold-standard immunohistochemical labeling with gel-assisted mass spectrometry imaging to enable cell-type-specific, single-cell spatial lipidomics with modern instrumentation. Using intact brain tissue as a testbed, we demonstrate in situ measurement of cerebellar Purkinje cells and spatially mapped lipids across timepoints and cerebellar subregions within the pathological landscape of a neurodegenerative, lysosomal storage disorder. With single-cell lipidomic profiling, we delineated spatiotemporally distinct accumulation of specific glycosphingolipids and phospholipids within Purkinje cells and non-Purkinje cells in the diseased brain. Furthermore, unsupervised single-cell lipidomic clustering elucidated disease-progression-and subregion-dependent molecular divergence between healthy and neurodegenerative states within the cerebellum.

## Main

Lipid homeostasis is fundamental to physiological function; the comprehensive characterization of lipid profiles and spatial organization thus provides critical mechanistic insights into healthy states and pathology, such as neurodegeneration^1^, metabolic disorders^2^, and oncogenesis^3–6^. Spatial lipidomics has emerged as a transformative modality for label-free spatial analysis of lipidomes within intact specimens, allowing visualization of localized lipid accumulation and depletion driven by lipid dyshomeostasis. To date, such spatial lipidomic studies have routinely relied on mass spectrometry imaging (MSI)^7^, a label-free analytical technique that combines mass spectrometry with spatial scanning. A highly versatile platform, MSI-based spatial lipidomics can be further integrated with other spatial omics modalities to reveal a broader and deeper molecular landscape of the specimen^8–11^. However, physical and instrumental resolution limits, particularly those inherent to matrix-assisted laser desorption/ionization mass spectrometry imaging (MALDI-MSI), have made it challenging for label-free spatial lipidomics to achieve single-cell or subcellular resolution across intact tissues using current instrumentation^12^.

Recent advances in sample expansion coupled with MALDI-MSI, including Gel-Assisted Mass Spectrometry Imaging (GAMSI)^13^ and other pioneering works^14–18^, have circumvented traditional resolution constraints of MALDI-MSI by physically magnifying the specimen while preserving its lipidomes or proteomes. However, the concurrent retention and visualization of lipid and protein information within a single specimen remains a key technical challenge. In GAMSI specifically, the use of fresh-frozen sample formats often results in the compromise of epitopes and cellular ultrastructure, while the conventional immunostaining process introduces a risk of lipid delocalization and loss. These hurdles constitute a critical methodological barrier to resolving the lipidomic profiles of individual cells identified by specific lineage or functional markers.

To overcome this challenge, we report immunohistochemistry-compatible GAMSI (or iGAMSI), a multimodal imaging platform allowing the concurrent spatial mapping of lipid and protein information within the same intact tissue specimen at the single-cell resolution. The iGAMSI workflow integrates fluorescence-based immunohistochemical labeling with gel-assisted MALDI-MSI, thereby eliminating the need to modify existing immunostaining protocols, mass spectrometry hardware, and analysis pipelines. Using brain tissue from a neurodegenerative disorder as proof-of-concept, we demonstrate that iGAMSI enables cell-type-specific, single-cell spatial lipidomic profiling of healthy and diseased states across complex tissue architectures.

## Results

### Development of iGAMSI and validation using intact tissue slices

We developed iGAMSI, a correlative fluorescence microscopy and MALDI-MSI workflow, to enable concurrent preservation of tissue ultrastructure, antigenic epitopes, and endogenous lipidome (**Fig. 1a, experimental procedure**). By utilizing chemical fixation and surfactant-free immunofluorescence (IF), iGAMSI ensures that the cellular lipid distribution remains largely unperturbed while allowing high-resolution protein and morphological imaging within intact tissues. In the iGAMSI workflow, an intact tissue slice (∼25 µm) undergoes pre-MSI fluorescence imaging after fixation and immunostaining to capture the spatial distribution of protein markers labeled via IF. The tissue slice is then polymerized in situ to form a superabsorbent hydrogel composite, homogenized by a controlled proteolytic process, and expanded multiple-fold linearly. After an immobilization and drying step, the sample-hydrogel composite is subject to MSI and post-MSI fluorescence imaging. In conjunction with its experimental workflow, we developed a post-processing pipeline for iGAMSI to perform pixel-level image registration of pre-MSI fluorescence, post-MSI fluorescence, and MS images (**Fig. 1a, post-processing and analysis, fig. S1**). Using the co-registered fluorescence and MS images, mass spectral profiles and their corresponding proteomic signatures can be extracted from individual cells, thereby enabling cell-type-specific spatial lipidomic mapping at the single-cell level.

**Fig. 1:**
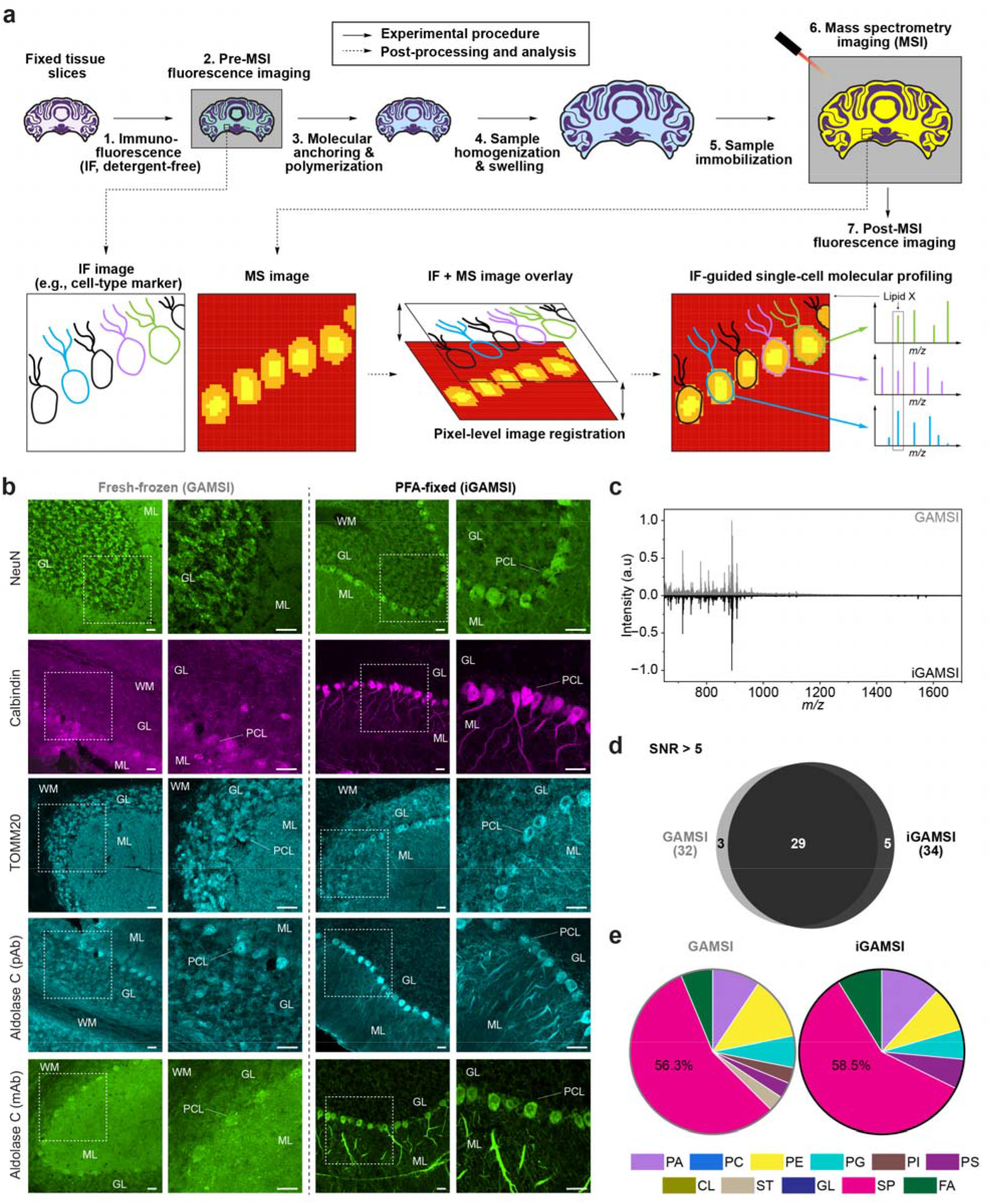
Immunochemistry-compatible gel-assisted mass spectrometry imaging (iGAMSI). a, Schematics showing the principle and generalized workflow of iGAMSI. Fixed tissue slices are (1) immunostained with fluorescently labeled antibodies (surfactant-free), (2) fluorescently imaged pre-mass spectrometry imaging (pre-MSI), (3) chemically anchored and polymerized to form a superabsorbent sample-hydrogel composite, (4) homogenized and expanded, (5) immobilized onto a sample plate or glass slide, (6) coated with matrix and analyzed on a mass spectrometer, and (7) fluorescently imaged post-MSI. The pre-MSI immunofluorescence (IF) image, post-MSI IF image, and mass spectrometry (MS) image are then co-registered to extract the single-cell mass spectra. **b**, Fluorescence images of fresh-frozen (left, two columns, as in the GAMSI workflow) and perfusion-fixed (right, two columns, as in the iGAMSI workflow) mouse cerebellum slices, immunostained for selected protein targets of (top to bottom) NeuN, calbindin, TOMM20, aldolase C (polyclonal; pAb), and aldolase C (monoclonal; mAb). Scale bars, 25 µm. **c,** Averaged mass spectra (*m/z* = 650-1700) of GAMSI (grey, top) and iGAMSI (black, bottom)-processed mouse cerebellum. Mass spectrum intensities were normalized to the base peak. **d,** Venn diagram showing lipid peaks identified from the averaged mass spectra of GAMSI (grey) and iGAMSI (black)-processed mouse cerebellum. All lipid assignments were made by comparing the lipid peaks with a >5 signal-to-noise ratio (SNR) to the LIPID MAPS database with a criteria of [M-H]^-^ and an allowed mass tolerance of *m/z* = ±0.1. **e,** Pie chart showing the chemical composition of GAMSI and iGAMSI lipid peaks in **d**. Instrument pixel size was set at 50 µm. PA: phosphatidic acid; PC: phosphocholine; PE: phosphoethanolamine; PG: phosphoglycerol; PI: phosphoinositol; PS: phosphoserine; CL: cardiolipin; ST: sterol lipid; GL: glycerolipid; SP: sphingolipid; FA: fatty acyl.

The successful implementation of iGAMSI required the optimization of a number of key steps in its workflow, such as the tissue fixation and homogenization conditions. In conventional MALDI-MSI techniques and the previously established GAMSI^13^, antigenic epitopes and cellular architectures can be compromised by the use of fresh-frozen tissue format^19,20^. Here, we performed paraformaldehyde (PFA)-based perfusion tissue fixation for iGAMSI, a fixation method preferred for immunohistochemistry (IHC) and known for superior preservation of tissue ultrastructure. In addition, PFA-fixed tissue has been previously validated to afford a lipidomic profile comparable to its fresh-frozen counterpart^21,22^. To evaluate antigenic and morphological preservation, we performed IF on both the fresh-frozen (as used for GAMSI) and PFA-fixed (as used for iGAMSI) mouse brain slices, targeting multiple common protein markers (**Fig. 1b**). The iGAMSI brain slices showed universal improvements in the quality of the immunostaining and the preservation of tissue ultrastructure, regardless of the antibody type and clonality tested. For instance, the cerebellar Purkinje cell (PC) dendrites commonly lost in the GAMSI brain slices were clearly preserved and visible with the iGAMSI method (**Fig. 1b**).

Another critical step within the iGAMSI workflow is the sample homogenization step, which proteolytically breaks down the structural integrity of the polymerized tissue while maintaining its native spatial and molecular information. Unlike GAMSI, which utilizes trypsin-based proteolysis to enhance lipid retention^13^, the stronger chemical fixation in iGAMSI necessitated a more rigorous homogenization for sample expansion. Consequently, we developed a surfactant-free, proteinase K (proK)-based proteolysis step for the iGAMSI workflow. In particular, the proteolysis kinetics and the composition of the homogenization buffer were systematically optimized to achieve a balance between isotropic, crack-free expansion, and maximal retention of lipids (**Table S3**).

To evaluate the retention of lipids, we collected mass spectra from iGAMSI and GAMSI-processed mouse cerebellum slices in both negative and positive modes. Qualitatively, we found that the iGAMSI samples replicated most of the major lipid ions (650-1700 *m/z* range) as observed from GAMSI, consistent with the previous reports supporting PFA-fixed samples for MALDI-MSI studies (**Fig. 1c**)^13^. To investigate the chemical compositions of the detected ion signals, we performed a library search and tabulated lipid peaks commonly detected across both samples, as well as those unique to either method (**Fig. 1d**, **fig. S2, fig. S3**)^23^. As a result, we found GAMSI and iGAMSI detected a comparable number of lipids. For instance, in the negative mode, the GAMSI and iGAMSI workflows resulted in 32 and 34 lipid assignments with a signal-to-noise ratio (SNR) of > 5, respectively, in one of the trials (**Fig. 1d**). In the positive mode, the GAMSI workflow resulted in 68 lipid assignments compared to 70 lipid assignments for the iGAMSI workflow with SNR > 5 in one trial (**fig. S3**). An assessment of the chemical distribution of the detected lipids demonstrated that major lipid classes were mostly conserved across GAMSI and iGAMSI, with primary-amine-containing lipids, including phosphatidylethanolamine (PE) and phosphatidylserine (PS), remaining largely unaffected (**Fig. 1e**). Collectively, these results demonstrate the efficacy of iGAMSI for in situ lipidomic profiling within intact tissue slices.

### iGAMSI enables cell-type-specific, single-cell lipidomic profiling across intact tissue

To demonstrate IF-guided single-cell spatial lipidomics, we applied iGAMSI to a coronal slice of wildtype (*Npc1^+/+^*, hereafter WT) mouse cerebellum. For initial validation, the tissue was immunostained against calbindin and aldolase C (or zebrin-II), two well-studied protein markers for cerebellar PCs^24–26^. Calbindin is a calcium-binding protein expressed in all PCs, and aldolase C is an enzyme expressed in a subset of PCs known for its role in glycolysis and neuroprotection^25,27,28^. We note that specific immunomarkers for the internal granular layer (IGL) were omitted, as the resident cell population consists of >99% granule cells^29^. These cells possess distinctive nuclear morphologies that are readily resolvable through the DAPI nuclear staining, permitting unambiguous identification without auxiliary labeling.

The application of the iGAMSI workflow allowed the acquisition of a spatially resolved MS image of the cerebellar tissue slice at an effective pixel size of ∼3.75 µm (15 µm instrument pixel size divided by the ∼4-fold sample linear expansion factor), a pixel size optimized to balance the SNR of MALDI-MSI with precise PC and non-PC registration (**Fig. 2a**). After the aforementioned image registration (**Fig. 1a, post-processing and analysis, fig. S1**), the IF-MS image overlay provided spatially-resolved lipidomic maps along with the protein marker information (**Fig. 2b**). The final overall registration accuracy was ∼3.75 µm, which was dictated by the effective pixel size of the MS images. This high accuracy was critical for the registration of PC and non-PC cell bodies (**Methods**). Notably, this single-cell registration was unattainable in the absence of gel-assisted tissue expansion with the native instrument pixel size of MSI (**fig. S4**).

**Fig. 2:**
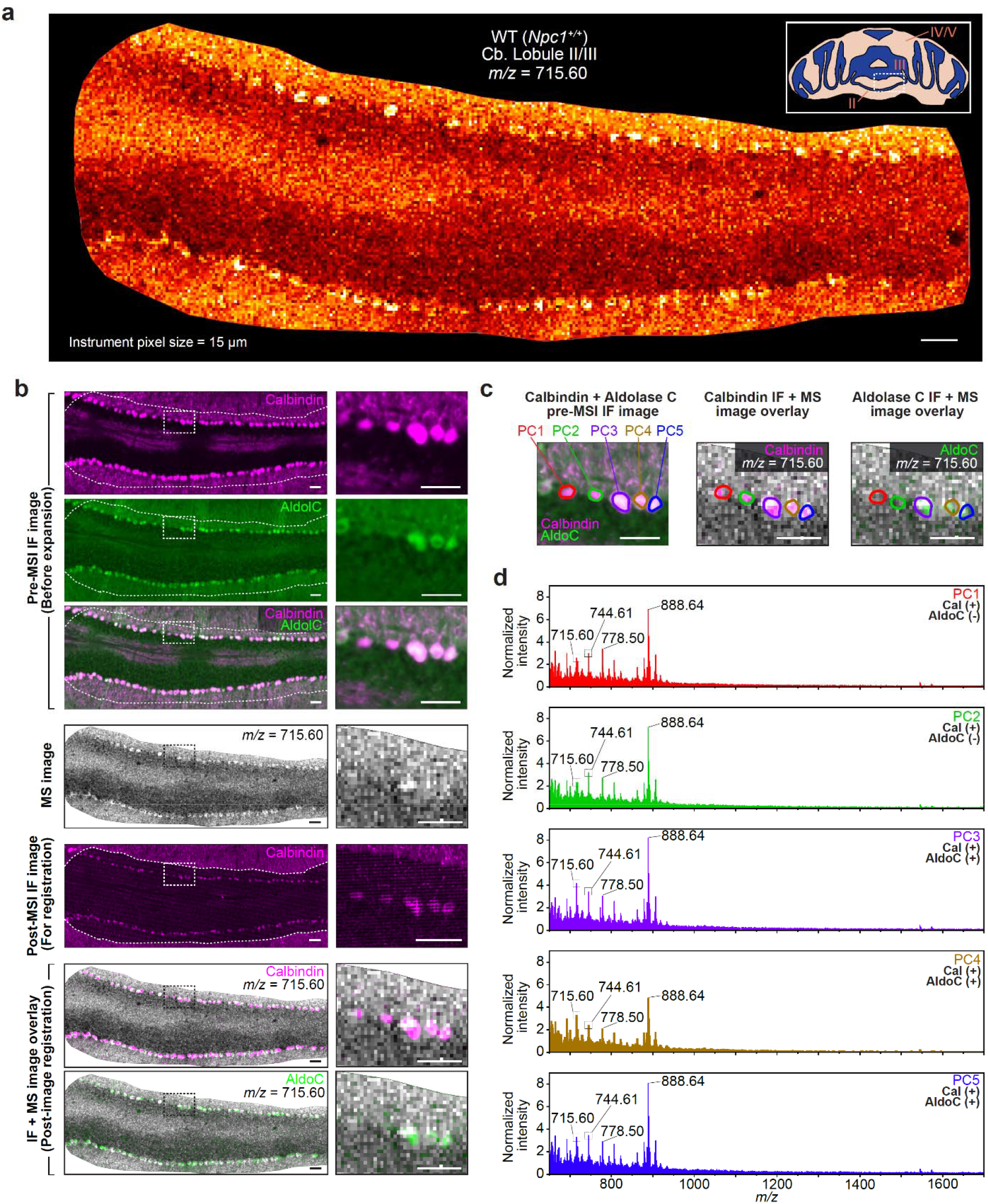
iGAMSI enables cell-type-specific single-cell lipidomic profiling. **a**, Spatial distribution of a selected mass spectrum peak (*m/z* = 715.60) in lobule III of an iGAMSI-processed wild-type (WT, *Npc1^+/+^*) mouse cerebellum slice. Inset, schematic illustration of a coronal mouse cerebellum slice, with white dotted box representing the MS image area. MSI experiments were performed on a rapifleX MALDI Tissuetyper (Bruker) with an instrument pixel size of 15 µm. Scale bar, 50 µm (200 µm). Here and after, unless noted otherwise, scale bars are provided at the pre-expansion scale with the post-expansion size shown in brackets. **b,** (Left, top to bottom) Pre-MSI IF images of calbindin (magenta), aldolase C (aldoC, green), calbindin and aldolase C overlay, MS image of a selected peak (*m/z* = 715.60; grey), post-MSI IF image of calbindin, registered pre-MSI calbindin IF and MS image overlay, and registered pre-MSI aldolase C IF and MS image overlay of the same iGAMSI-processed WT mouse cerebellum slice in **a**. Scale bars, 50 µm (200 µm). (Right, top to bottom) Magnified views of the outlined regions on the left column. Scale bars, 50 µm (200 µm). **c,** Magnified views of (left to right) pre-MSI calbindin (magenta) and aldolase C (green) IF image, registered pre-MSI calbindin IF and MS (*m/z* = 715.60; grey) image overlay, and registered pre-MSI aldolase C IF and MS image overlay of the outlined regions in **b**, showing individually segmented Purkinje cells (PCs). **d,** Single-cell mass spectra (*m/z* = 650-1700) of individual PCs from **c**. Scale bars, 50 µm (200 µm). cal (+): calbindin-positive; cal (-): calbindin-negative; aldoC (+): aldolase-C-positive; aldoC (-): aldolase-C-negative.

Next, using the cell-type protein markers as guidance, we extracted the lipidomic profiles of multiple neighboring PCs (**Fig. 2c**). Mass spectra of each PC (650-1700 *m/z* range) revealed that the same ion (*m/z* = 715.60) was consistently enriched within PCs across the same cerebellar lobule (**Fig. 2d**), while control spectra from the hydrogel alone confirmed minimal background signal contribution (**Fig. S5)**. Collectively, these results validate that the iGAMSI workflow can serve as a reliable platform for cell-type-specific, single-cell MALDI-MSI analyses of lipids and other molecular features within complex tissue architectures.

### iGAMSI reveals accumulation of key glycosphingolipids in cerebellar PCs in a model of neurodegeneration

Niemann-Pick disease, Type C (NPC) is a progressive and neurodegenerative disorder caused by NPC1 or NPC2 gene mutations, which disrupt lysosomal transport and lead to the pathological accumulation of cholesterol and glycosphingolipids^30^. These mutation-specific molecular signatures trigger severe neurodegeneration across multiple brain regions, including the cerebellum, thalamus, and cortex, resulting in the decline and loss of motor and cognitive functions among other symptoms^31,32^.

Previous studies using mutant NPC mouse models found that gangliosides GM1 (d36:1), GM2 (d36:1), and GM3 (d36:1) are the key storage glycosphingolipids involved in NPC progression^33,34^. Specifically, detailed immunocytochemistry studies revealed pronounced GM2 accumulation within PCs in the null mutant NPC1 mouse (*Npc1^-/-^*, hereafter NPC1) cerebella^35^. However, the abundance of GM1 and GM3 could not be fully validated using this immunostaining-based approach^36^. In recent MALDI-MSI time-course studies, most of the cerebellar regions with abundant GM2 and GM3 presence showed less elevation in GM1, consistent with previous reports of GM2 and GM3 accumulation resulting from GM1 degradation^34,37^. Interestingly, in the posterior and flocculonodular lobules of the NPC1 cerebellum, with a particular pronounced increase in lobule X, all three gangliosides were found elevated at the pre-terminal age (9 weeks). However, the spatial resolution limitation of these MALDI-MSI studies precluded cellular-level investigation of such localized accumulation. This left the elevation or depletion of these gangliosides in specific cell types, particularly within cerebellar PCs, unvalidated at the pre-terminal stage using MALDI-MSI.

To elucidate the distribution of key gangliosides GM1, GM2, and GM3 within PCs in the pre-terminal NPC1 mouse cerebellum, we performed cell-type-specific iGAMSI on 9-week-old WT and NPC1 mouse cerebellum focusing on lobule X (**Fig. 3a-3c**). Here, lobule X was selected for our initial validation for two reasons: the aforementioned observation of enhanced ganglioside localization in this lobule, and prior studies indicating it is the only region to maintain a substantial number of surviving PCs at this disease stage^37,38^ (**Fig. 3a**). Notably, we found pronounced elevation of all three gangliosides, GM1, GM2, and GM3, within PCs of lobule X in the NPC1 cerebellum (**Fig. 3b-3c**). In contrast, no elevation was observed within the WT PCs of lobule X (**Fig. 3b-3c**). To further validate this genotypic enrichment, we plotted the average mass spectra (650-1700 *m/z* range) of twenty randomly selected PCs from each genotype (**Fig. 3d**). Consistent with the MS images, we confirmed that GM1, GM2, and GM3 were elevated in the NPC1 PCs as compared to the WT PCs.

**Fig. 3:**
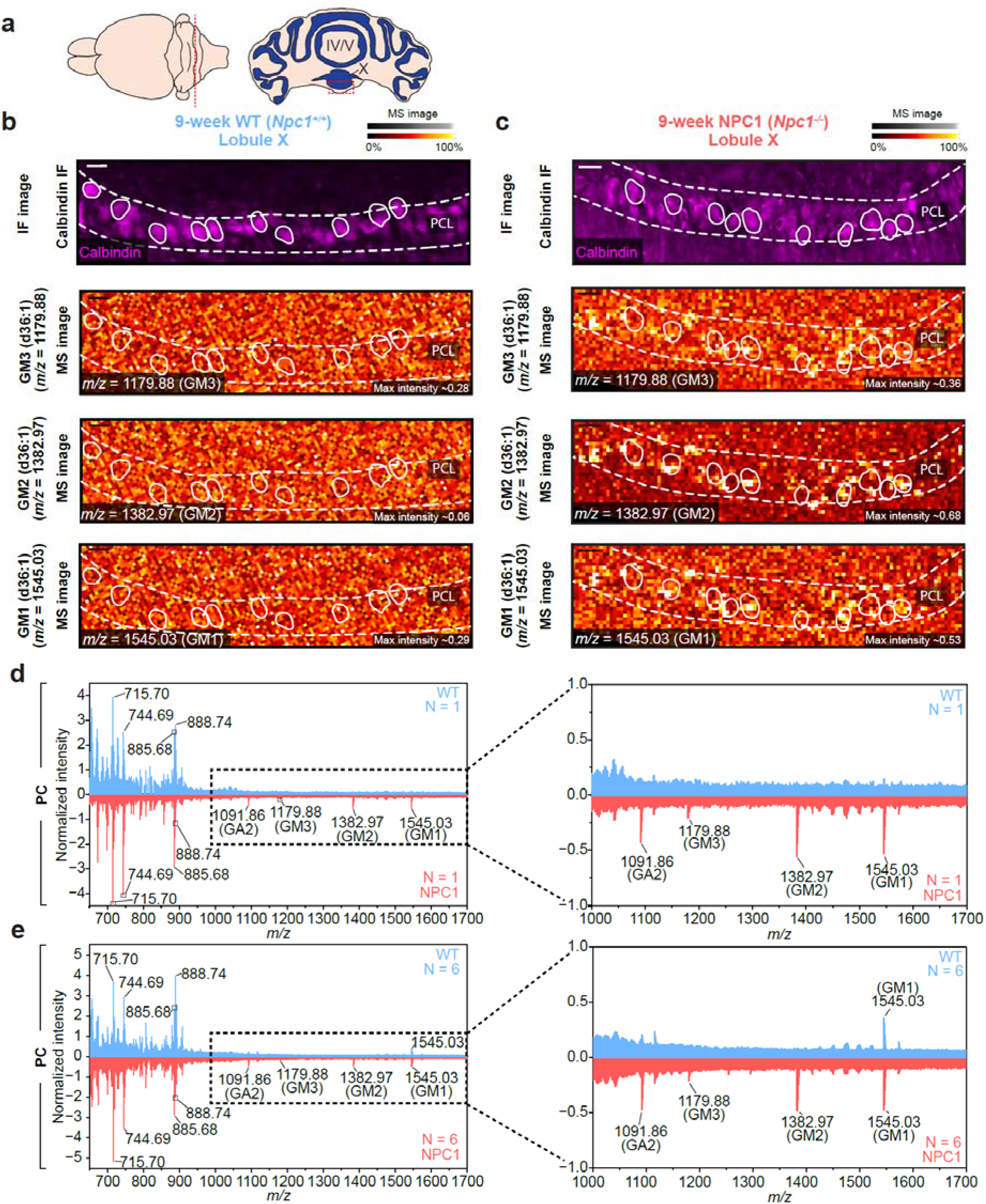
Cell-type-specific accumulation of sphingolipids within cerebellar PCs. a, Schematic illustration of (left) the mouse brain and (right) a coronal mouse cerebellum slice, with the red dashed line representing the coronal section and the red dashed box representing the IF and MS image area. **b-c,** (Top to bottom) calbindin IF images (magenta), MS image of ganglioside GM3 (d36:1) (*m/z* = 1179.88; red hot), MS image of ganglioside GM2 (d36:1) (*m/z* = 1382.97; red hot), MS image of ganglioside GM1 (d36:1) (*m/z* = 1545.03; red hot) within a part of lobule X (red dashed box in **a**) of 9-week (**b**) Wild-type (WT, *Npc1^+/+^*) and (**c**) Niemann-Pick disease type C1 (NPC1, *Npc1^-/-^*) mouse cerebellum. Scale bars, 25 µm (100 µm). Several PCs are outlined by white circles in the MS images as an example. Max intensity values within the MS images indicate the highest pixel values across the respective imaged areas. **d,** (Left) Averaged mass spectra (*m/z* = 650-1700) of PCs in the lobule X of the 9-week WT (blue) and NPC1 (red) mouse cerebellum in **b**-**c** (biological replicate N = 1). (Right) Magnified views of the dashed box region in the left plot. **e,** (Left) Averaged mass spectra (*m/z* = 650-1700) of PCs in the lobule X of 9-week WT (blue) and NPC1 (red) mouse cerebella (N = 6; 3 males and 3 females). (Right) Magnified views of the dashed box region in the left plot.

To account for biological variability, we further performed iGAMSI-based single-cell lipidomic profiling across six animals (three males and three females). As a result, we found similar elevations of GM1, GM2, and GM3 in the NPC1 PCs of lobule X across all animals (**fig. S6 and S7**). No sex-based differences were detected. The combined average mass spectra of PCs further confirmed the elevation of all three gangliosides in the NPC1 PCs compared to the WT PCs (**Fig. 3e**). Interestingly, asialo-GM2 (d36:1) (GA2) (*m/z* = 1091.86) also displayed elevated levels in the NPC1 sample (**fig. S8**). While GA2 is a recognized metabolic byproduct of GM2 degradation^39^, our findings provide the first evidence of a concurrent elevation of GM2 and GA2 within the pre-terminal PCs in NPC1 cerebellum. We note that GM1 was also found elevated in the averaged WT mass spectrum, but was not consistently detected in all WT mice. These results confirm a genotypic elevation of GM1, GM2, GM3, and GA2 within PCs of NPC1 animals, suggesting a key metabolic role for these glycosphingolipids in PC survival and degeneration at the disease’s terminal stage.

### iGAMSI elucidates spatiotemporally distinct accumulation of sphingolipids and phospholipids in cerebellar PCs and non-PCs

To assess lipidomic alterations associated with disease progression, we quantified the abundance of the aforementioned key glycosphingolipids within PCs of lobule X in both pre-symptomatic (4-week) and pre-terminal (9-week) NPC1 mouse cerebella. In agreement with our mass spectrometry analyses and previous reports, we found a statistically significant increase in GM1, GM2, GM3, and GA2 within NPC1 PCs of lobule X across the two time points (**Fig. 4a**)^35,37^. In contrast, for non-PCs, which are predominantly granule cells, we observed non-significant changes of these ganglioside species except for GM3, which showed a significant increase in the NPC1 samples (**Fig. S9**). Next, we examined the abundance of these glycosphingolipids within both aldolase-C-positive and negative PCs of lobule IV/V in the pre-symptomatic mouse cerebella (**Fig. 4b**). Interestingly, our results showed no statistically significant differences between the two PC subtypes, regardless of their genotypes. However, a significant genotypic difference was observed for GA2 and GM2, indicating that ganglioside sequestration manifests as early as the pre-symptomatic stage (4 weeks) within both aldolase-C-positive or negative PCs in the NPC1 mice (**Fig. 4b**). These results also suggest that, contrary to our initial hypothesis, the sequestration of these glycosphingolipids may not be directly correlated with the expression of aldolase C within PCs.

**Fig. 4:**
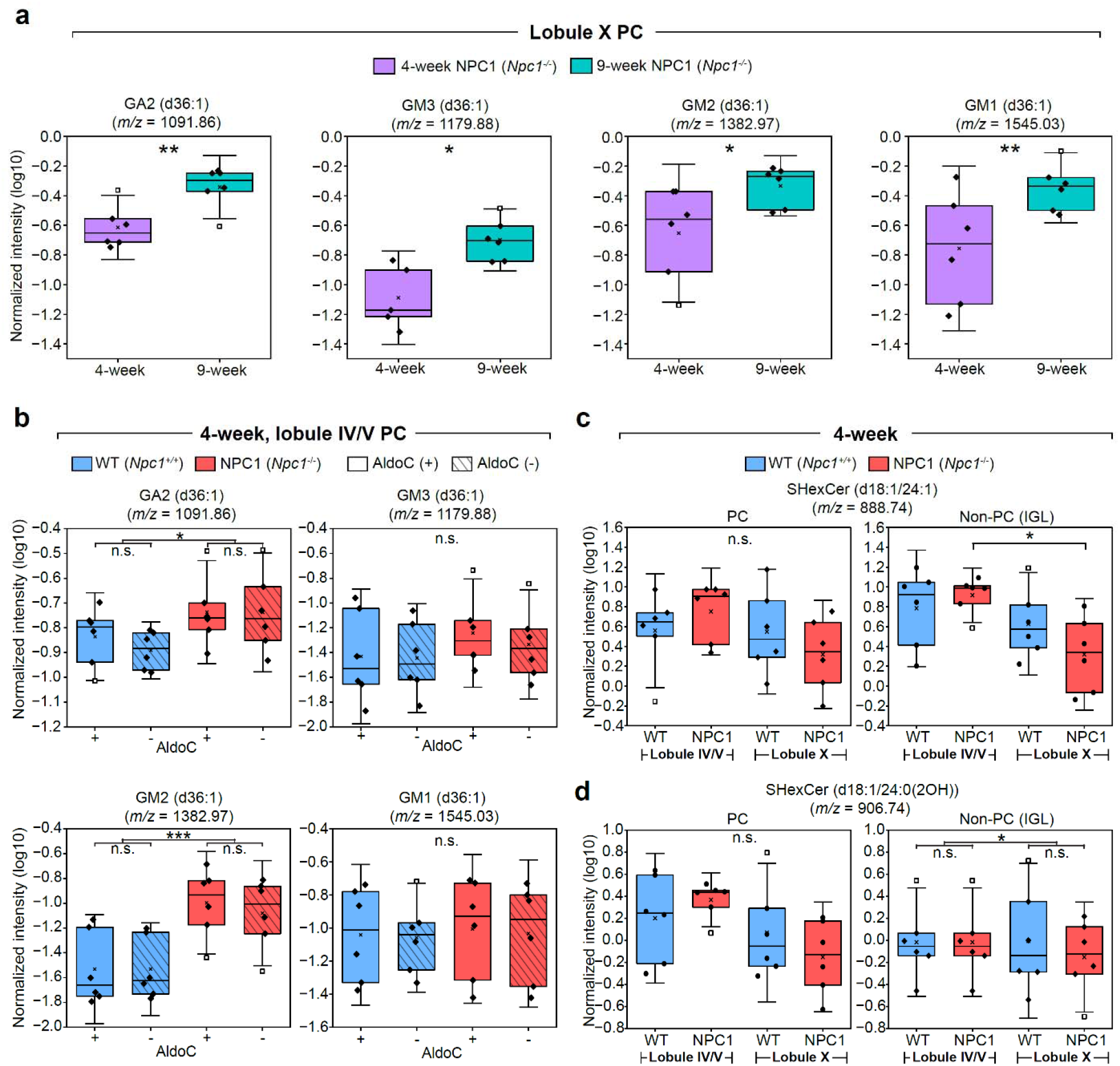
Disease-progression-and lobule-dependent accumulation of lipids in cerebellar PCs and non-PCs. a, (Left to right) Relative abundance of GA2 d36:1 (*m/z* = 1091.86), GM3 d36:1 (*m/z* = 1179.88), GM2 d36:1 (*m/z* = 1382.97), and GM1 d36:1 (*m/z* = 1545.03) in PCs of lobule X in 4-week (purple) and 9-week (teal) NPC1 mouse cerebella. The *p* values for the unpaired t-tests (with Welch’s correction) were *p* = 0.008 (GA2), 0.0096 (GM3), 0.055 (GM2), and 0.043 (GM1). **b,** (Top to bottom, left to right) Relative abundance of GA2, GM3, GM2, and GM1 in aldolase-C-positive (solid) and negative (striped) PCs of lobule IV/V in 4-week WT (blue) and NPC1 (red) mouse cerebella. A 2-way ANOVA with Tukey’s test was performed with *p* values of *p* = 0.025 (GA2), and 0.0003 (GM2) (with the complete set of *p* values provided in **Table S4**). We note that in the 4-week GM3 plot in a and b, only five data points are shown as the log transform of one data point was an undefinable value. **c-d,** Relative abundance of (**c**) SHexCer (d18:1/24:1) (*m/z* = 888.74) and (**d**) SHexCer (d18:1/24:0(2OH)) (*m/*z = 906.74) in (left) PCs and (right) non-PCs (within the inner granule layer, IGL) of lobules IV/V and X in 4-week WT (blue) and NPC1 (red) mouse cerebella. A 2-way ANOVA followed by Tukey’s test was performed with *p* values of *p* = 0.027 (for non-PCs in **c**) and 0.030 (for non-PCs in **d**) (with the complete set of *p* values provided in **Tables S5** and **Table S6**). All the relative abundance of lipids are represented as box plots, where the ends of the whiskers represent standard deviation +/-1.5 times the interquartile range, the upper line of the box represents the 75^th^ percentile, the middle line represents the 50^th^ percentile (median), the lower line represents the 25^th^ percentile, the diagonal cross represents the mean, and the open squares represent the individual values of the outlying data points. *: *p* < 0.05; **: *p* < 0.01; ***: *p* < 0.001; ****: *p* < 0.0001; n.s.: not significant (*p* ≥ 0.05).

Previous reports have indicated that lipid dysregulation occurs in a spatially heterogeneous manner across the cerebellum during NPC1 progression, especially among different lobules^33,37^. To characterize this spatial heterogeneity, we quantitatively compared the lipidomic profile of PCs and non-PCs across the anterior (IV/V) and the flocculonodular (X) lobules. Interestingly, a subclass of sphingolipids, sulfatides (SHexCer), showed significant variances between these lobules in the pre-symptomatic NPC1 cerebella (**Fig. 4c-4d, fig. S10**). For instance, SHexCer (d18:1/24:1) (*m/z* = 888.74) exhibited a statistically significant difference between lobules IV/V and X for NPC1 non-PCs (**Fig. 4c**). We note that while no statistically significant differences were observed for this lipid within PCs, its abundance followed a similar but nonsignificant downward trend for NPC1 PCs of lobule X. Similarly, SHexCer (d18:1/24:0(2OH)) (*m/z* = 906.74) showed a statistically significant decrease in lobule X (**Fig. 4d**) across both genotypes in non-PCs. Another structurally similar sulfatide SHexCer (d18:1/22:0(2OH)) (*m/z* = 878.73), however, exhibited no significant differences at the lobule or genotype level, but showed a nonsignificant downward trend within the NPC1 non-PCs of lobule X (**fig. S11a)**. Finally, phosphoinositide PI (18:0/20:4) (*m/z* = 885.68) has shown a downward trend in lobule IV/V as well (**fig. S11b)**. This result corroborates prior studies reporting a decreased level of this particular PI in NPC1 cerebella through disease progression^40^. Our study further indicates that this decreased expression begins as early as 4-weeks within the granule cell layer in a lobule-specific manner. Together, these results suggest that previously unknown lipid alterations occur within PCs and non-PCs in a highly lobule-dependent manner within the NPC1 cerebella.

### iGAMSI enables single-cell lipidomic clustering and delineation of wild-type and neurodegenerative cell populations

The datasets generated by the iGAMSI workflow are inherently high-dimensional, necessitating application of additional bioinformatic analyses to extract and interpret biologically meaningful patterns. To reduce the data dimensionality, we applied Uniform Manifold Approximation and Projection (UMAP) to the single-cell lipidomic data. After validating and applying a batch correction pipeline, the single-cell lipidomic UMAP enabled visualization of latent structures within complex lipidomic profiles that were challenging to delineate with lipid-specific analyses (**Methods, Supplementary Notes 1**).

Using this single-cell lipidomic clustering pipeline, we investigated the molecular alterations across the NPC1 disease progression in a cell-type-specific manner. We first performed analysis on PCs and found clear genotype-dependent clustering from the 9-week lobule X PCs (**Fig. 5a-5c)**. Marker analysis revealed GA2, GM1, GM2, PI (18:0/20:4), and an unknown feature of *m/z* = 857.57 were the top 5 features distinguishing the clusters (**fig. S12**). In comparison, 4-week PCs of lobule IV/V did not show clear genotypic clustering, suggesting that PC lipidomic profiles do not differ substantially across WT and NPC1 in lobule IV/V at this stage of disease. Additionally, no difference in the clustering pattern was observed between 4-week aldolase-C-positive and negative PCs of lobule IV/V (**Fig. 5a** and **fig. S13**). This further confirmed that the PC lipidomic profiles are similar across the aldolase-C subtypes.

**Fig. 5:**
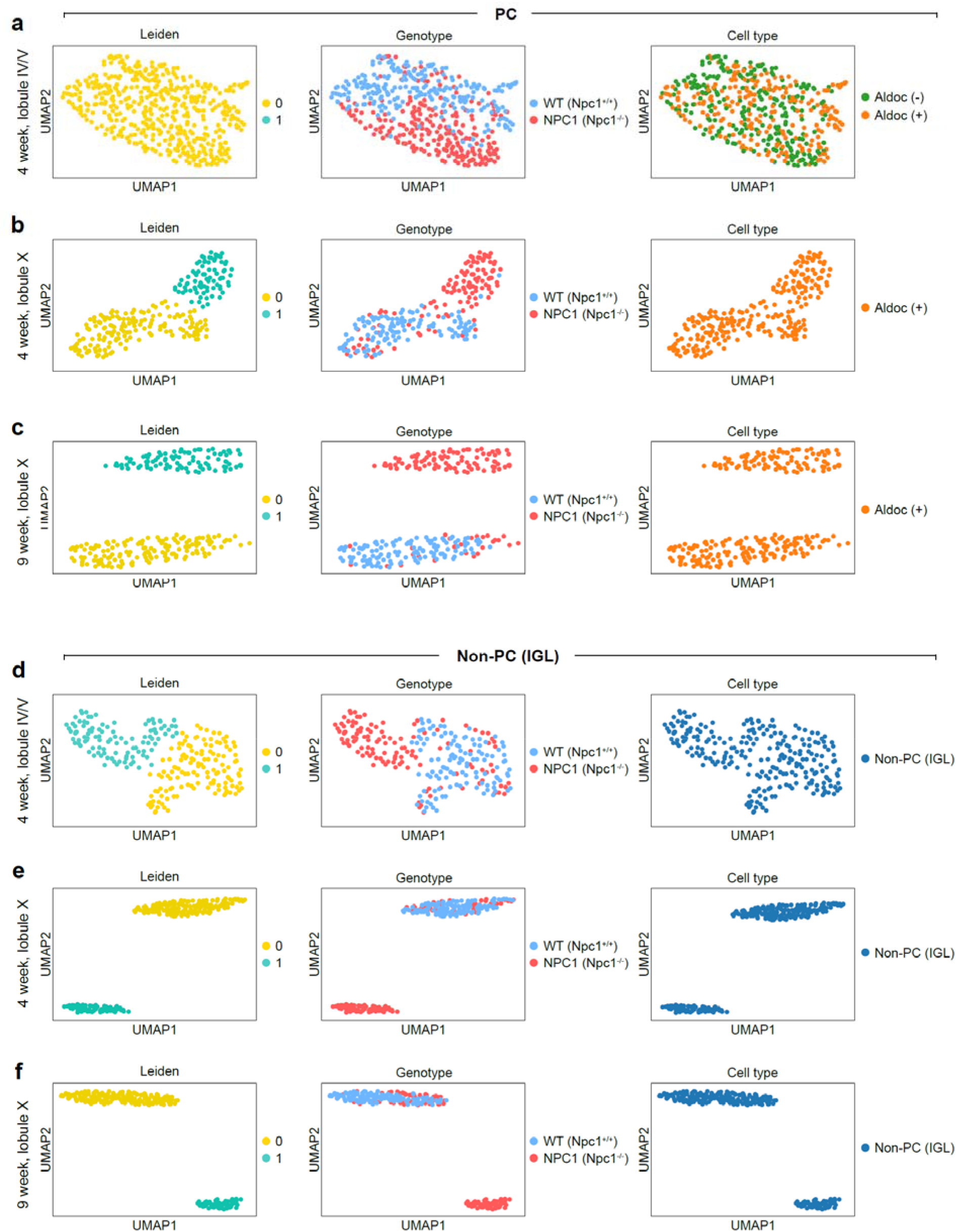
iGAMSI enables single-cell lipidomic clustering. a-c, UMAP embedding of PCs of (**a**) lobule IV/V (4-week), (**b**) lobule X (4-week), and (**c**) lobule X (9-week) cerebella, annotated by (left to right) Leiden clusters, genotypes, and cell types. **d-f,** UMAP embedding of non-PCs (IGL) from lobule IV/V in (**e**) lobule IV/V (4-week). (**f**) lobule X (4-week), and (**g**) lobule X (9-week) cerebella, annotated by (left to right) Leiden clusters, genotypes, and cell types. All UMAP embeddings were computed from neighbor graphs constructed from the data, with the Leiden clustering performed at a resolution of 1.0. AldoC (+): aldolase-C-positive; AldoC (-): aldolase-C-negative. The cluster indices (0, 1,…,n) correspond to the first, second,…,and (n+1)th clusters generated by Leiden clustering.

For non-PCs, a different clustering pattern emerged. In particular, genotypic clustering was found for both 4-and 9-week-old mice, with increased separation in lobule X (**Fig. 5d-5f**). For 4-week non-PCs of lobule IV/V, the top 5 lipids driving the clustering were SHexCer (d18:1/24:1), *m/z* = 656.29, SHexCer (d18:1/22:0(2OH)), *m/z* = 862.60, and 890.63 (**fig. S14a**). For 4-week non-PCs of lobule X, the top 5 lipids driving the clustering were SHexCer (d18:1/24:1), *m/z* = 890.63, PE (18:0/22:6), PI (18:0/20:4), and *m/z* = 657.30 (**fig. S14b**). Markedly, 9-week non-PCs of lobule X mice showed similar differential lipid features as the 4-week non-PCs of lobule X, with SHexCer (d18:1/24:1), *m/z* = 890.63, PE (18:0/22:6), PI (18:0/20:4), and *m/z* = 656.29 as the top 5 lipids (**fig. S14c**). These results confirmed a lobule-specific difference in the lipidomic profile of non-PCs at the early disease stage, with lobule X non-PCs showing a more distinctive lipidomic difference compared to lobule IV. Moreover, the lipids driving genotypic clustering for non-PCs were distinct from those driving the PCs, highlighting the cell-type-specific nature of lipidomic alterations across the disease states.

## Discussion

Using a multimodal imaging pipeline combining fluorescence and high-spatial-resolution mass spectrometry imaging, iGAMSI allows single-cell spatial mapping of lipids and protein labels within the same intact tissue. We validated the utility of iGAMSI by characterizing the lipidomic profiles of cerebellar PCs and non-PCs in brain tissue slices from a mouse model of neurodegenerative lysosomal storage disorder NPC1. Our method revealed a distinct spatiotemporal pattern of lipid accumulation, comprising a distinct array of sphingolipids and phospholipids, both previously known but understudied, across the cerebellar lobules and along the trajectory of disease progression.

Previous studies found aldolase-C-negative PCs are more susceptible to neurodegeneration in NPC1 mouse^38^. As the anterior lobules are populated with a greater number of aldolase-C-negative PCs, the PC degeneration appears to proceed with an anterior to posterior spatial pattern as the disease progresses. A recent lipid MSI study showed a similarly patterned accumulation of gangliosides GM2 and GM3 in the posterior lobules along with the disease progression^37^. However, whether such accumulation occurs within the degenerating PCs or other cellular structures has remained unknown. Our findings thus provide evidence of a concurrent elevation in GM1, GM2, and GM3 within the persistent Purkinje cell populations of the posterior lobules, highlighting a specific lipidomic resilience or vulnerability of aldolase-C-positive PCs. We also found that GA2, a previously understudied glycosphingolipid, also follows a similar pattern of accumulation in PCs^41^. However, contrary to our initial hypothesis, the lipidomic profiles of aldolase-C-positive and negative PCs did not differ significantly at the pre-degeneration stage. This suggests that the neuroprotective nature of aldolase-C expression may be driven by other metabolic determinants or that there could be other molecular pathways for PC cytotoxicity in NPC1^42^.

Sulfatides are a structural glycolipid crucial in maintaining the myelin sheaths along with differentiation of oligodendrocytes, and their loss has been associated with NPC1 primarily through myelin-specific lipidomics studies^43–45^. The significant decrease of multiple SHexCer in PCs and non-PCs of lobule X in our study is one of the first evidence that this subclass of lipids is also altered in the neuronal cells in the NPC1 brain. Specifically, our results suggest that across multiple neuronal cell types in the internal granular layer, sulfide dysregulation could occur as early as 4-weeks in a lobule-specific manner. While future study is required to identify the specific metabolic pathways involving these sulfatides in neuronal cells, such cell-type-specific and spatial effects could not be delineated from bulk or multi-cell lipidomics data.

Aside from cell-type-specific lipidomic profiling, our method, for the first time, enabled unsupervised single-cell lipidomic clustering from intact, complex tissue architecture^46,47^. Our clustering results indicate that genotypic clustering of PC lipidomic profiles is both disease-progression and lobule-specific, with gangliosides being the dominant driver for the clustering. In contrast, sulfatides and PIs are the main drivers in the genotypic clustering of non-PCs. These results highlight the importance of spatially-resolved single-cell lipidomic analyses in understanding the spatiotemporal heterogeneity in cellular lipidome in complex tissue architecture.

From a methodological perspective, further enhancements in the achievable spatial resolution are possible by increasing the expansion factor. For the iGAMSI workflow reported here, the expansion factor of ∼4-fold was optimized to achieve an effective lateral resolution of <5 µm with commercially available mass spectrometers, which ensures the spatial registration errors remain below the single-cell threshold while maintaining analytically efficient signal counts and acquisition times. By introducing high-expansion-factor hydrogel chemistry (e.g., with ∼10-20-fold expansion), the effective pixel size of iGAMSI can potentially be reduced to the nanoscopic length scale for subcellular spatial lipidomics studies. With robust preservation of cellular ultrastructure and epitopes, spatial lipidomic mapping of organelles, neuronal processes, and synapses might be possible with iGAMSI.

An inherent constraint of our methodology is the reduction in analyte density following sample expansion, as lipids (and other molecules) are diluted in proportion to the square of the expansion factor. This sensitivity trade-off may be mitigated by ongoing advancements in the detection limits of next-generation mass spectrometers. In addition, the mass spectra from the current implementation of iGAMSI are typically convolved with hydrogel background peaks in the *m/z* < 650 range, complicating the extraction of lipidomic features, including those for cholesterols. Further experimental optimizations as well as computational advances to reduce and deconvolve the hydrogel background may mitigate such limitations. Moreover, the management of overlaid multimodal datasets presents a primary computational hurdle. The escalating scale and dimensionality of these data necessitate AI-driven frameworks for the automated segmentation of cellular masks and the integration of multi-omic molecular signatures ^48^. As the field of single-cell spatial lipidomics matures, we anticipate the emergence of standardized bioinformatic pipelines to streamline these automated workflows.

Despite these constraints, iGAMSI circumvents current technical barriers to provide an integrated modality for cell-type-specific and functionally annotated single-cell spatial lipidomics using conventional and best-practice immunohistochemistry and mass spectrometry. Because iGAMSI requires no specialized hardware modifications or analysis pipelines, it remains accessible to the existing research facility with standard mass spectrometry infrastructure. Leveraging its high-throughput and multimodal imaging capability, iGAMSI is poised to generate the extensive datasets required to construct cell-type-informed lipidomic foundation models, offering deeper understandings of metabolic diversity across intact tissues in healthy and diseased states at the single-cell level.

## Methods

### Ethics statement

The animal care and handling procedures, including basic care, housing and husbandry, and surgical procedures, were performed in accordance with the US National Institutes of Health Guide for the Care and Use of Laboratory Animals and approved by the University of Illinois Chicago Animal Care Committee.

### Chemicals and reagents

All reagents were used as supplied unless noted otherwise. Purified water was obtained from a Milli-Q IQ 7000 Ultrapure Water System (MilliporeSigma). All chemicals and reagents were obtained from MilliporeSigma unless noted otherwise. Detailed antibody information is provided in **Supplementary Tables 1 and 2**.

### Animals

Adult, both male and female, heterozygous BALB/cNctr-*Npc1*^m1N^/J mice (Jackson Laboratory) were maintained in a breeding colony at the University of Illinois Chicago. All animals were housed in appropriate cages with 12-hour dark and 12-hour light cycles, ambient temperature, and humidity. Genotyping was performed using polymerase chain reaction as previously described^35,37^. The primer sequences used were (FWD8F) 5′-GGTGCTGGACAGCCAAGTA-3′ and (REVINTR3) 5′-GATGGTCTGTTCTCCCATG-3′.

### Tissue preparation

Wild-type control (*Npc1^+/+^*) and NPC1 mutant (*Npc1^−/−^*) mice at 4 and 9 weeks of age were anesthetized using 5% isoflurane (SomnoFlo Low-Flow Electronic Vaporizer, Kent Scientific) and transcardially perfused with 30□mL of 1x PBS and then 30 mL of 4% paraformaldehyde (PFA) (047340.9M, ThermoFisher) in 1x PBS. All solutions were maintained at a temperature of 4 °C. After perfusion, the brains were dissected, post-fixed in 4% PFA in 1x PBS at 4 °C for ∼20-24□hours, and stored in 1x PBS at 4 °C. The fixed brains were sectioned using a vibratome (VT1200S, Leica) to ∼25 μm coronal or sagittal slices and stored in 1x PBS at 4 °C in a 24-well plate until further procedures.

### Immunohistochemistry

For immunofluorescence, the brain slices were incubated in a detergent-free blocking buffer [5% (v/v) normal goat serum (NGS) (005-000-121, Jackson Immuno Research) in 1x PBS] at room temperature for 1 hour, and then in a 300 nM DAPI (1023627600, MilliporeSigma) solution at room temperature for 1 hour. Next, the brain slices were incubated in the primary antibody solution (1:100 dilution with the detergent-free blocking buffer) shaking overnight at room temperature. The brain slices were then washed four times with the blocking buffer for 30 min each time, and incubated in the secondary antibody solution (1:200 dilution with the detergent-free blocking buffer) shaking overnight at room temperature. Finally, the brain slices were washed four times with 1x PBS (or the detergent-free blocking buffer) for 30 min each time, and stored in 1x PBS at 4 °C until further procedures.

The stored brain slices were carefully transferred using a paintbrush and mounted on positively charged microscope slides (MID7100-45, MIDSCI) after immunostaining. Briefly, the microscope slides were first treated with poly-L-lysine solution [0.1 % (w/v) in water] (P8920-100ML, MilliporeSigma) for 15 min and washed with water twice for 15 min each time. The brain slices were then carefully transferred to and mounted on these pre-treated microscope slides using a paintbrush.

### Pre-mass spectrometry imaging (MSI) fluorescence imaging

Pre-MSI fluorescence images of the immunostained brain slices were obtained using a Nikon spinning disk confocal system (CSU-W1, Yokogawa) with a 10x 0.45 NA air objective (Nikon) in wide-field mode. Unless noted otherwise, all fluorescence microscopy data in this work were collected using NIS Elements AR v5.30.04 (Nikon).

### Sample gelation

The immunostained brain slices were incubated in an acryloyl-X, SE (AcX, A20770, ThermoFisher) solution (0.1 mg/mL in 1x PBS) at room temperature for 6 hours to overnight. The brain slices were then washed twice with 1x PBS for 15 min each time. Gelation of the brain slices was performed following a modified version of previously described protocols^29,30^. Briefly, the gelation chamber was created with 25 μm tape surrounding the mounted brain slices. The brain slices were then incubated with ∼50 μL of gelation solution [1x PBS, 2 M NaCl, 8.625% (w/v) sodium acrylate, 2.5% (w/v) acrylamide, 0.15% (w/v) N,N’-methylenebisacrylamide, 0.01% (w/v) of 4-hydroxy-2,2,6,6-tetramethylpiperidin-1-oxyl (4HT), 0.2% (w/v) of ammonium persulfate (APS), and 0.2% (w/v) of tetramethylethylenediamine (TEMED)] with a #1.5 coverglass of placed atop the solution and gelled in a humidified 37 °C incubator for 2 hours.

### Sample homogenization, expansion, and immobilization

The gelled brain slices were trimmed into a right trapezoid to track the orientation of the sample and homogenized in a detergent-free proteinase K (ProK) (P8107S, New England Biolabs) homogenization buffer (8 units/mL in 1x PBS) at room temperature for 4 hours.

The gelled and homogenized brain slices were examined, and if necessary, flipped to ensure the tissue side faced upwards. The gelled brain slices were then placed in an excess volume of 0.5x PBS for 20 min and then in purified water three times for 20 min each time until the samples were fully expanded.

The expanded samples were transferred onto the ITO-coated glass slides (CB-90IN-S111, Delta technologies) or stainless steel MALDI plates (Applied Biosystems SCIEX) using a 36 x 60 mm no. 1.5 cover glass (260461-100, Ted Pella), ensuring the tissue side is facing upwards. Finally, the samples were dried under vacuum in a Pyrex vacuum desiccator filled with Drierite at room temperature for 4 hours to overnight.

### Matrix application

1,5-diaminonaphthalene (DAN) was applied to the immobilized and dried sample via sublimation using a previously described homemade sublimation apparatus^38,39^. Briefly, 50 mg of DAN was dissolved in 2 mL of acetone, aspirated onto the bottom of the sublimation flask, and blow-dried using nitrogen to form a thin layer of white solid. Next, a hotplate, to which the sublimation flask was placed, was set to 105 °C while a digital thermometer was placed in contact with the bottom of the flask to monitor the temperature. An ice slush was added to the cold finger of the apparatus, to which the ITO-coated glass slide or the MALDI plate was adhered on the underside with copper tape. Finally, the sublimation apparatus was placed under vacuum at 80 mTorr using a rough pump, and the DAN matrix was sublimed for 2 min. The amount of matrix deposited on the sample was determined as mass per square centimeter.

### Mass spectrometry imaging (MSI)

Mass spectrometry imaging was performed using a rapifleX MALDI Tissuetyper equipped with a Smartbeam 3D 10 kHz Nd:YAG (355 nm) laser (Bruker). The rapifleX MALDI Tissuetyper was operated in negative or positive ion reflection mode to acquire data between mass ranges (*m/z*) of 500-2000 using DAN as the matrix. Mass spectrometry (MS) images were acquired with either 15 µm or 50 µm raster distance and 200 laser shots per pixel using flexControl 4.0 (Build 46, Bruker). The laser intensity was adjusted to maintain an optimal signal intensity (e.g., ∼1 x 10^4^).

### Post-MSI fluorescence imaging

Post-MSI fluorescence images of the brain slices were obtained using a Nikon spinning disk confocal system (CSU-W1, Yokogawa) with a 10x 0.45 NA air objective (Nikon) in wide-field mode.

### Image registration

Images from the fluorescence imaging and MSI were registered in two steps. First, non-rigid registration was performed between the post-MSI fluorescence image and the MS image using the BigWarp 9.0.0 plugin on ImageJ distribution Fiji (ver. 1.53t). The post-MSI fluorescence image was set as the moving image and registered to the MS image by aligning the laser ablation marks in the fluorescence images with the pixels in the MS image as the fiducial markers. Next, non-rigid registration was performed between the pre-and post-MSI fluorescence images using the same BigWarp plugin. The pre-MSI fluorescence image was set as the moving image and registered to the post-MSI fluorescence image using the PC somas as the registration landmarks. Finally, the registered pre-MSI fluorescence image and the MS image were overlaid using ImageJ.

## Data processing

All MS data processing, including region-of-interest (ROI) definition, average mass spectrum extraction, and image generation, was performed on SCiLS Lab (ver. 2024a, Bruker) using the overlaid pre-MSI fluorescence images as a reference, unless noted otherwise. Briefly, all raw MS data from the rapifleX MALDI Tissuetyper (Bruker) instrument were imported in SciLS lab using linear binning with a bin size of 16.018 mDa and a mass range of *m/z* 500-2000. For single-cell analysis, separate ROIs corresponding to individual cells were created using the overlaid pre-MSI fluorescence image. Average mass spectra from each ROI were subsequently exported for further analyses.

For Venn diagram generation, the averaged mass spectra across the ROIs are exported to the mMass software (available at: https://github.com/xxao/mMass-Dist/tree/main/v5.5.0)^23^. Deisotoping was performed using the following parameters: maximum charge = 1, isotope mass tolerance = 0.1 m/z, isotope intensity tolerance = 50%, isotope mass shift = 0, label envelope = ‘1st selected’, envelope intensity = ‘envelope maximum’, with the options ‘Remove isotope’, ‘Remove unknown’, and ‘Set labels as monoisotopes’ enabled. Peak picking was performed with the following settings: S/N threshold of 5, Abs. intensity threshold of 0, Rel. intensity threshold of 0%, picking height of 50. Baseline correction was enabled (precision = 15; relative offset = 25), followed by Savitzky–Golay smoothing (window size = 0.3 m/z; 2 cycles). Deisotoping was applied during peak picking, and shoulder peaks were removed. Lipid assignments were made by comparing the experimental mass measurements to the LIPID MAPS database (www.lipidmaps.org) with an allowed mass tolerance of *m/z* = ±0.1. When multiple lipid candidates matched the same experimental mass, assignments were evaluated based on mass error and the carbon side-chain composition of the candidates.

For cell-type-specific single-cell lipidomic analyses, the extracted mass spectra of individual cells from the same cell type and age were averaged using simple averaging. Then the data were centroided, aligned, and normalized to the internal standard PE (18:0/22:6) (*m/z* = 790.66) using a custom code modified from the MALDIquant pipeline^49^. The PE peak was selected as the internal standard because prior study found that it stays statistically unchanged in abundance across different genotypes throughout the disease progression^43^. In detail, the MS data were first preprocessed using Savitzky-Golay smoothing with a half window size of 5. Then, baseline removal was performed using the SNIP option with an iteration of 100. Next, the data were centroided using the ‘detectPeak’ function with ‘SuperSmoother’, a half window size of 10, and SNR of 1. After centroiding, the mass spectra were aligned with a mass tolerance of ±0.15. Intensities of the mass spectra were then normalized to the intensity of the internal standard PE (18:0/22:6) (*m/z* = 790.66) peak within each mass spectrum.

Unless noted otherwise, all MS images were exported using Total Ion Count (TIC) on SCiLS lab. The *m/*z values for ion images depicted are based on the centroided and aligned data. For the relative abundance plots, the y-axis represents the log_10_ ratio of the target lipid intensity normalized to the internal standard.

## Statistical analysis

For statistical analysis, data were analyzed by an unpaired t-test or a two-way or three-way ANOVA followed by a Tukey’s multiple comparisons post hoc test. Differences were considered statistically significant with a *p* value of <0.05.

### Clustering

For cellular clustering analysis, a master feature list was first generated. Briefly, mass spectra from the same cell type of the same animal were averaged, centroided, and aligned with SNR > 10. These feature lists were then combined into a master feature list by merging shared features while retaining all unique features observed in any dataset. Next, feature extraction was performed for individual cells by comparing the centroided peaks within each mass spectrum to the master feature list, with a tolerance of *m/z* = ±0.3. Next, batch correction was performed using ReCombat to remove the batch effects arising from the sample and instrumental variances^50^. We defined each sample run, which involves tissue fixation, immunostaining, gelation, expansion, matrix application, and MSI run without any sample exchanges, as a separate batch. Detailed parameters can be found in (https://github.com/HoraceChan99/NonRigidReg.git). Isotopic features were manually reviewed based on inspection of the corresponding MS data. Features that fall within the isotopic distribution of another peak are classified as isotopes and excluded from differential analysis. The Leiden clustering resolution used is indicated in each figure.

### GAMSI sample preparation

GAMSI samples were prepared as control following previously described protocols^13^. For immunohistochemistry validation, the cryo-sectioned brain slices were thawed to room temperature and briefly fixed with 4% PFA in 1x PBS at room temperature for 10 min. After removing the fixatives, the brain slices were washed with 1x PBS three times for 5 min each time, and immunostained using the same procedure as the iGAMSI protocol.

### Fluorescence imaging for immunohistochemistry validation

Fluorescence images of the immunostained brain slices were obtained using a Nikon spinning disk confocal system (CSU-W1, Yokogawa) with a 10x 0.45 NA air objective (Nikon), a 20x 0.95 NA water immersion objective, and a 40x 1.15 NA water immersion objective in wide-field and confocal modes.

### Tandem mass spectrometry (MS/MS)

MS/MS was performed with the 4800 Plus MALDI TOF/TOF Analyzer (Applied Biosystems SCIEX) in negative ion reflection mode using DAN as the matrix. The instrument was operated in the 2 kV, gas off, operation mode to allow unimolecular decay. Mass spectrometry data from 4800 Plus MALDI TOF/TOF Analyzer (Applied Biosystem SCIEX) were collected using 4000 Series Explorer v5.5.3 (Applied Biosystem SCIEX).

Additional MS/MS was performed with the timsTOF fleX QTOF Analyzer (Bruker) in negative ion mode using DAN as the matrix. The instrument was operated with the following parameters: CID: 40-70 eV; isolation width: 2.5 *m/z*; pre-pulse storage: 10 µs. Source conditions are as follows: nebulizer: 0.3 bar; dry gas: 3.5 V/min; dry temp: 200 °C.

### Statistics and reproducibility

No statistical method was used to predetermine sample size. Biological replicates of six animals of each genotype, three males and three females, were collected. No data was excluded from the analysis. The extraction of single-cell ROIs was randomized. The investigators were blinded to allocation during experiments and outcome assessment. For reproducibility, all the validation experiments were repeated independently at least three times using tissue from the same animal, unless noted otherwise.

## Data availability

The mass spectrometry data generated in this study have been deposited in the public MassIVE repository under accession code xxx [link]. Source data are provided with this paper.

## Code availability

Custom scripts used for data analysis are available on GitHub at https://github.com/HoraceChan99/NonRigidReg.git.

## Acknowledgments

We thank D. Pierre-Jacques and W. Li of the Cologna lab at the University of Illinois Chicago for their helpful discussions and support. We thank S. Shafaie, F. Tobias, B. Owen, and the Northwestern University IMSERC facility for assistance with mass spectrometry imaging. We thank Chicago Biomedical Consortium (CBC) for access to core mass spectrometry facilities. R.G. acknowledges funding support from US NIH DP2MH136390, US NIH UG3MH126864, Searle Scholars Program, McKnight Technological Innovations in Neuroscience Award, and University of Illinois Chicago Startup Fund. S.M.C. acknowledges funding support from US NIH R01NS114413, US NIH R01NS124784, and US NSF CAREER Award 2143920.

## Author contributions

Conceptualization: R.G. and S.M.C.

Methodology: M.C.H., Y.H.C., and R.G.

Investigation: M.C.H., Y.H.C, H.P.D., and K.C.P.

Formal analysis: M.C.H. and Y.H.C.

Software: Y.H.C. and M.C.H.

Visualization: M.C.H., Y.H.C., and R.G.

Funding acquisition: R.G. and S.M.C.

Project administration: R.G. and S.M.C.

Supervision: R.G. and S.M.C.

Writing – original draft: M.C.H., Y.H.C. and R.G.

Writing – review & editing: all authors.

