## Supplemental Information for "Immunohistochemistry-compatible gel-assisted mass spectrometry imaging"

**The PDF file includes:**

Supplementary Notes

Supplementary Figs. S1 to S17

Supplementary Tables S1 to S6

#### Supplementary Notes

##### Supplementary Note 1: Validation of batch correction methods in iGAMSI pipeline

Due to the limitation in the physical size of the expanded sample, imaging data were collected over multiple imaging sessions. Initial analysis revealed a pronounced batch effect arising from instrumental variability and differences in sample preparation. Direct clustering of normalized data yielded clusters formed primarily by imaging batch rather than biological condition (**fig. S15a**), highlighting the need for effective batch correction. We therefore evaluated two widely used approaches, ComBat and ReComBat[1–4], and selected ReComBat based on its performance in mitigating batch-associated variation (**fig. S15b-c**).

To assess whether ReComBat introduced overfitting or artificially enhanced biological separation, we designed controlled validation experiments. First, to test label-driven bias, we isolated 9-week wild-type samples and reassigned half of the labels to a pseudo-NPC1 group. Prior to batch correction, clustering reflected batch-driven separation. However, after ReComBat processing, the data collapsed into a single cluster (**fig. S16**), indicating that the algorithm does not impose artificial label-based structure. Second, we analyzed datasets comprising repeated measurements from the same NPC1 animal, along with data from two wild-type animals. Following ReComBat correction, samples were grouped by genotype, although not separated at the current clustering resolution (**fig. S17**), demonstrating that biological variation is preserved while technical variation is effectively reduced. Together, these results confirm that ReComBat enables reliable disentanglement of biological signals from technical noise without overfitting.

### Supplementary Figures

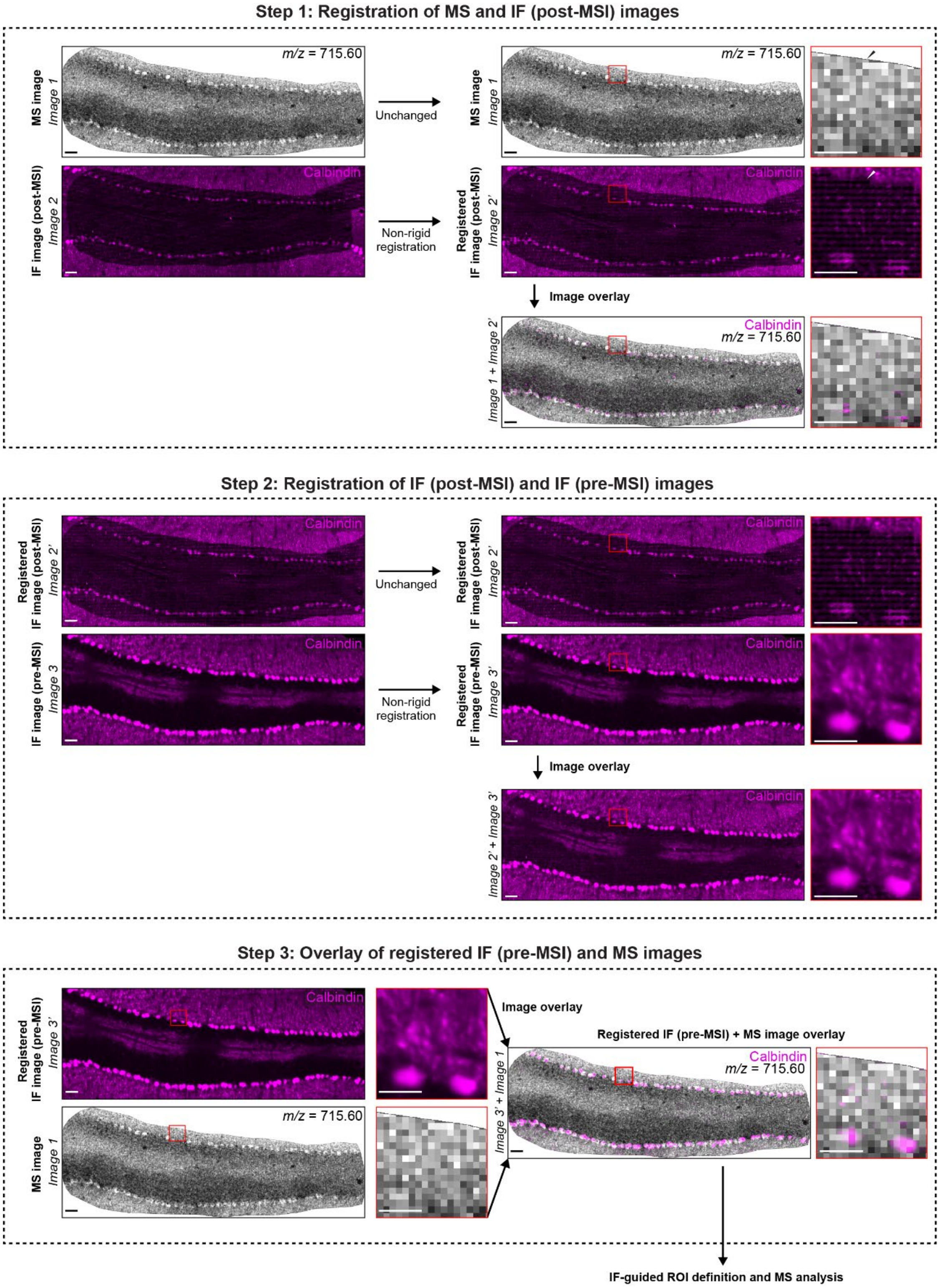

**Supplementary Fig. S1: iGAMSI image registration pipeline.** (Top) Step 1: Registration of MS and IF (post-MSI) images. Column 1: MS image (Image 1) of select lipid (*m/z* = 715.60) and IF image (post-MSI, Image 2) stained for calbindin (magenta). Image 1 remains unchanged while Image 2 is non-rigidly registered to MSI (Image 1) by using the laser ablation marks and pixels as the fiducial markers. Column 2: MS image (Image 1), Registered IF image (post-MSI, Image 2'), and Overlay of Image 1 and Image 2'. Column 3: Magnified views of the regions outlined in red boxes in previous images (left). Black/white arrow indicates the same pixel in each image. (Middle): Registration of IF (post-MSI) and IF (pre-MSI) images. Column 1: Registered IF image (post-MSI, Image 2') and IF image (pre-MSI, Image 3) stained for calbindin (magenta). Image 2' remains unchanged while Image 3 is non-rigidly registered to Image 2' using the Purkinje cells as the fiducial markers. Column 2: Registered IF image (post-MSI, Image 2'), Registered IF image (pre-MSI, Image 3'), and Overlay of Image 2' and Image 3'. Column 3: Magnified views of the regions outlined in red boxes in previous images (left). (Bottom) Step 3: Overlay of registered IF (pre-MSI) and MS images. Column 1: Registered IF image (pre-MSI, Image 3') and MS image (Image 1). Column 2: Magnified views of the regions outlined in red boxes in previous image (left). Column 3: Overlay of Image 3' and Image 1. Column 4: Magnified view of region outlined in red box in previous image. Mass spectrometry images were collected on a rapifleX MALDI Tissuetyper (Bruker) with an instrument pixel size of 15 µm. Scale bars, 50 µm (200 µm). For magnified images, scale bars, 25 µm (100 µm).

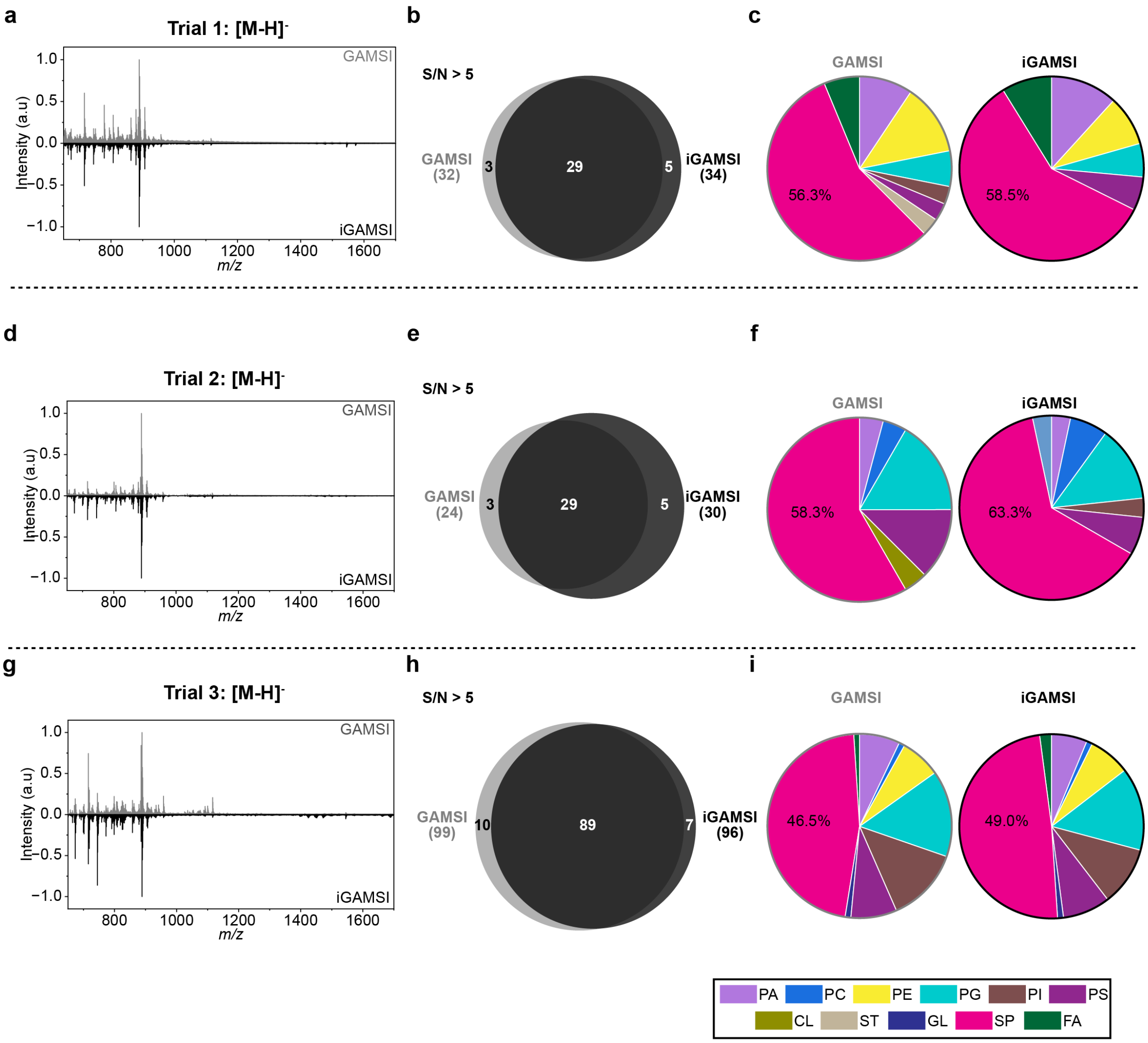


**Supplementary Fig. S2: Lipidomic profiles of GAMSI and iGAMSI-processed samples (trials 2 and 3, negative ion mode, [M-H]$^-$). a, d, g,** Averaged mass spectra (*m/z* = 650-1700) of GAMSI (grey, top) and iGAMSI (black, bottom)-processed mouse cerebellum. Mass spectrum intensities were normalized to the base peak. **b, e, h,** Venn diagram showing lipid peaks identified from the averaged mass spectra of GAMSI (grey) and iGAMSI (black)-processed mouse cerebellum. All lipid assignments were made by comparing the lipid peaks with a >5 signal-to-noise ratio (SNR) to the LIPID MAPS database with an allowed mass tolerance of *m/z* = ±0.1. **c, f, i,** Pie chart showing the chemical composition of GAMSI and iGAMSI lipid peaks in **d**. Instrument pixel size was set at 50 μm. PA: phosphatidic acid; PC: phosphocholine; PE: phosphoethanolamine; PG: phosphoglycerol; PI: phosphoinositol; PS: phosphoserine; CL: cardiolipin; ST: sterol lipid; GL: glycerolipid; SP: sphingolipid; FA: fatty acyl. Trial 1 corresponds to the data presented in **Fig. 1c-1e**.

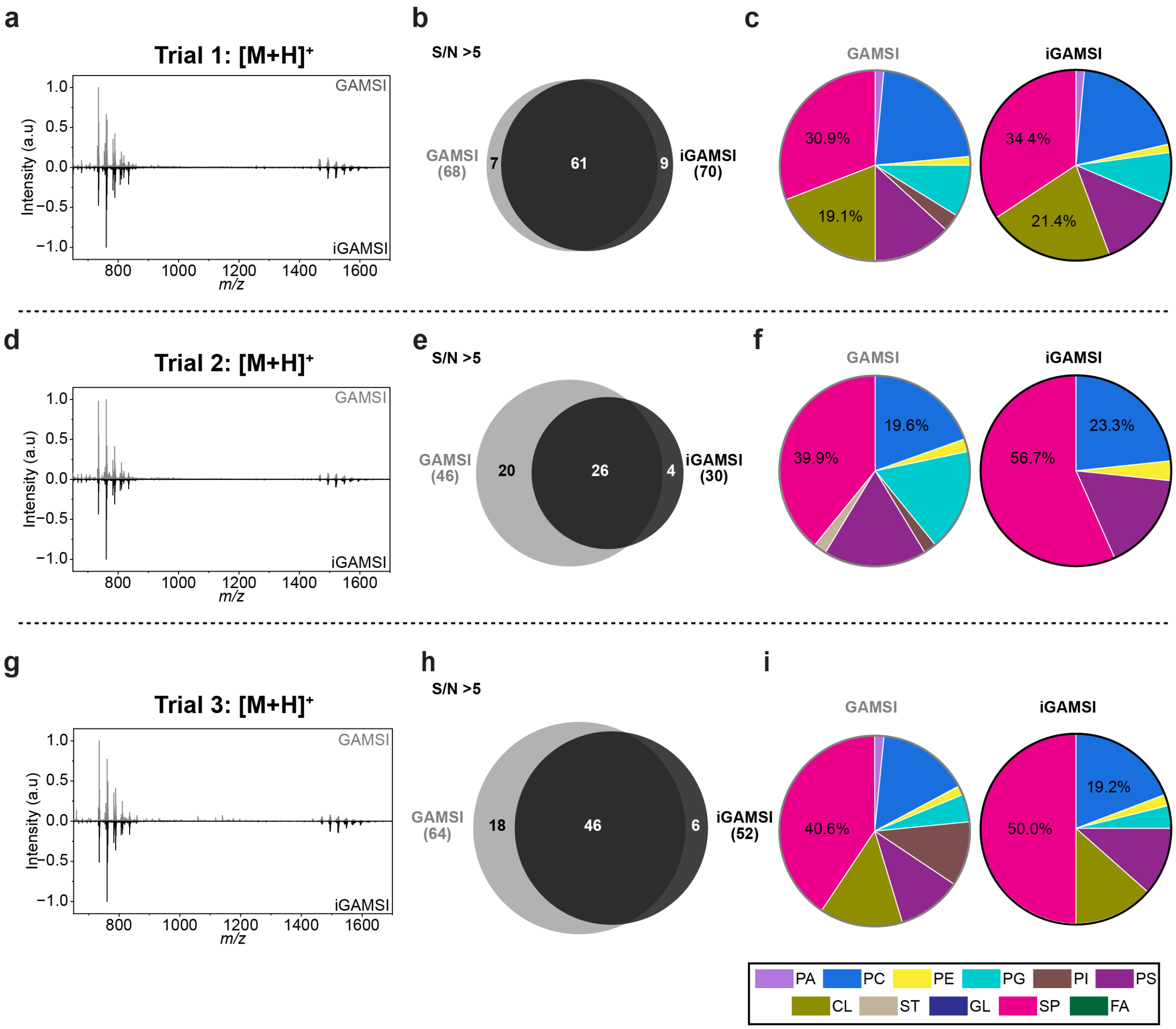


**Supplementary Fig. S3: Lipidomic profiles of GAMSI and iGAMSI-processed samples (positive ion mode, [M+H]$^+$). a, d, g,** Averaged mass spectra ($m/z$ = 650-1700) of GAMSI (grey, top) and iGAMSI (black, bottom)-processed mouse cerebellum. Mass spectrum intensities were normalized to the base peak. **b, e, h,** Venn diagram showing lipid peaks identified from the averaged mass spectra of GAMSI (grey) and iGAMSI (black)-processed mouse cerebellum. All lipid assignments were made by comparing the lipid peaks with a >5 SNR to the LIPID MAPS database with a criterion of [M+H]$^+$ and an allowed mass tolerance of $m/z$ = ±0.1. **c, f, i,** Pie chart showing the chemical composition of GAMSI and iGAMSI lipid peaks in **a, d, g**. Instrument pixel size was set at 50 µm. PA: phosphatidic acid; PC: phosphocholines; PE: phosphoethanolamine; PG: phosphoglycerol; PI: phosphoinositol; PS: phosphoserine, CL: cardiolipin; ST: sterol lipid; GL: glycerolipid; SP: sphingolipid; FA: fatty acyl**.**

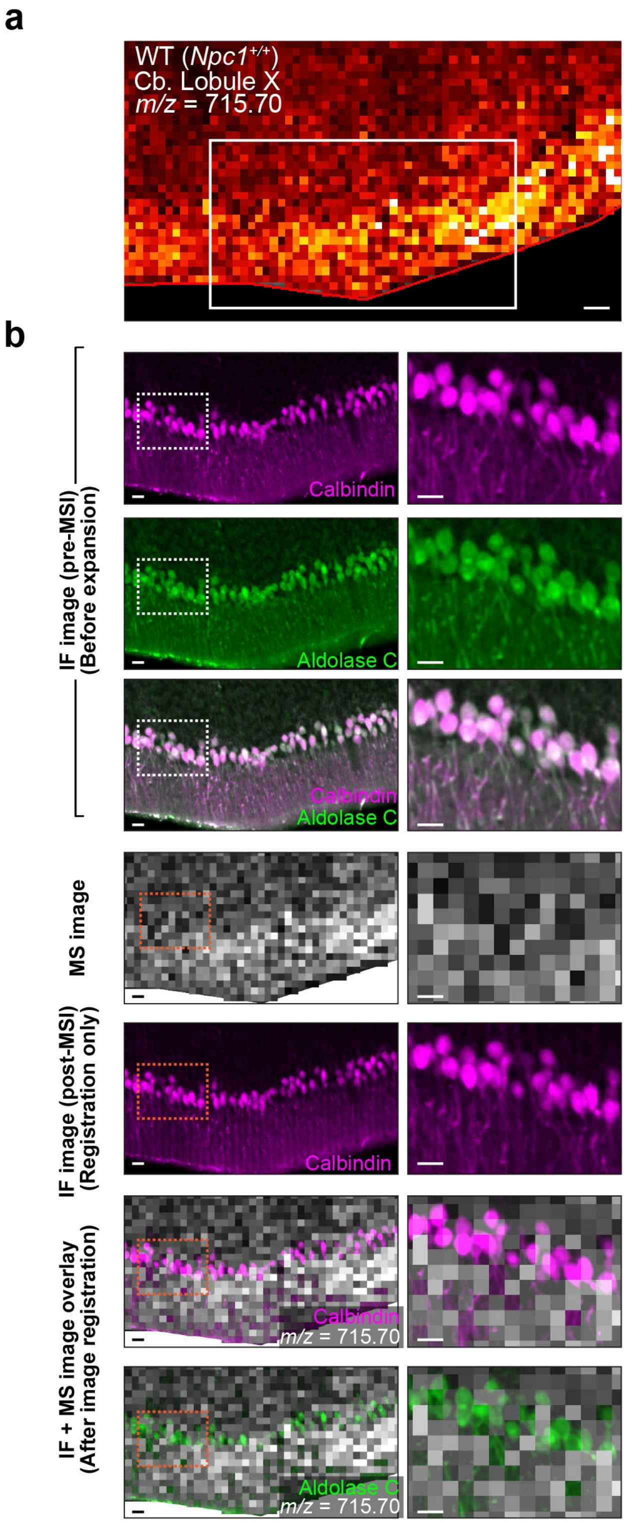
a
WT (Npc1+/+)
Cb. Lobule X
m/z = 715.70
b
IF image (pre-MSI)
(Before expansion)
Calbindin
Aldolase C
Calbindin
Aldolase C
MS image
IF image (post-MSI)
(Registration only)
Calbindin
IF + MS image overlay
(After image registration)
Calbindin
m/z = 715.70
Aldolase C
m/z = 715.70

**Supplementary Fig. S4: Multimodal image registration without gel-assisted sample expansion. a,** Spatial distribution of a selected mass spectrometry peak (*m/z* = 715.70) in lobule X of an unexpanded WT mouse cerebellum slice with the white solid box representing the zoomed in images in **b.** MSI experiments were performed on a rapifleX MALDI Tissuetyper (Bruker) with an instrument pixel size of 15 μm. Scale bar, 50 μm. **b,** (Left) Pre-MSI IF images of calbindin (magenta), aldolase C (green), calbindin and aldolase C overlay, MS image of a selected mass spectrometry peak (*m/z* = 715.70; grey), post-MSI IF image of calbindin, registered pre-MSI calbindin IF and MS image overlay, and registered pre-MSI aldolase C IF and MS image overlay of the same unexpanded WT mouse cerebellum slice in **a.** Scale bars, 25 μm. (Right) Magnified views of the outlined regions. Scale bars, 25 μm.

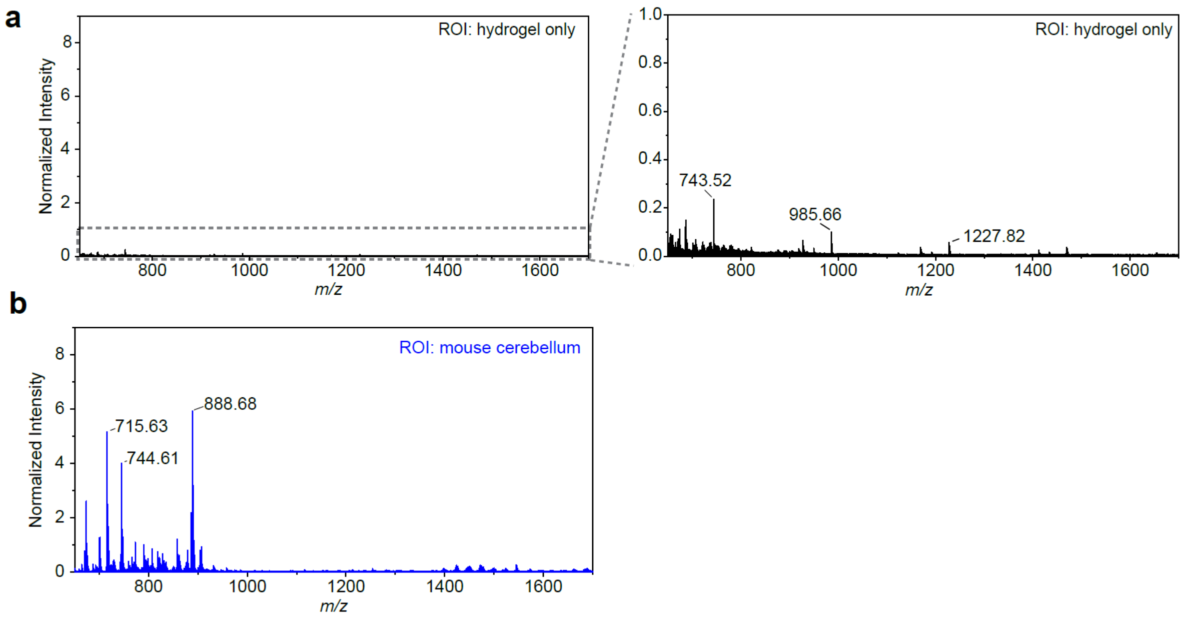


**Supplementary Fig. S5: Hydrogel background of iGAMSI-processed samples. a,** (left) Averaged mass spectra (*m/z* = 650-1700) of twenty regions-of-interest (ROIs) extracted from the hydrogel-only regions. (Right) Magnified view of the dashed box region in the left plot. **b,** Averaged mass spectra (*m/z* = 650-1700) of twenty ROIs extracted from the mouse cerebellum sample. Mass spectra were collected on a rapifleX MALDI Tissuetyper (Bruker) with an instrument pixel size of 15 µm. All spectra were centroided and normalized to the internal standard PE (18:0/22:6) (*m/z* = 790.66) in **b**.

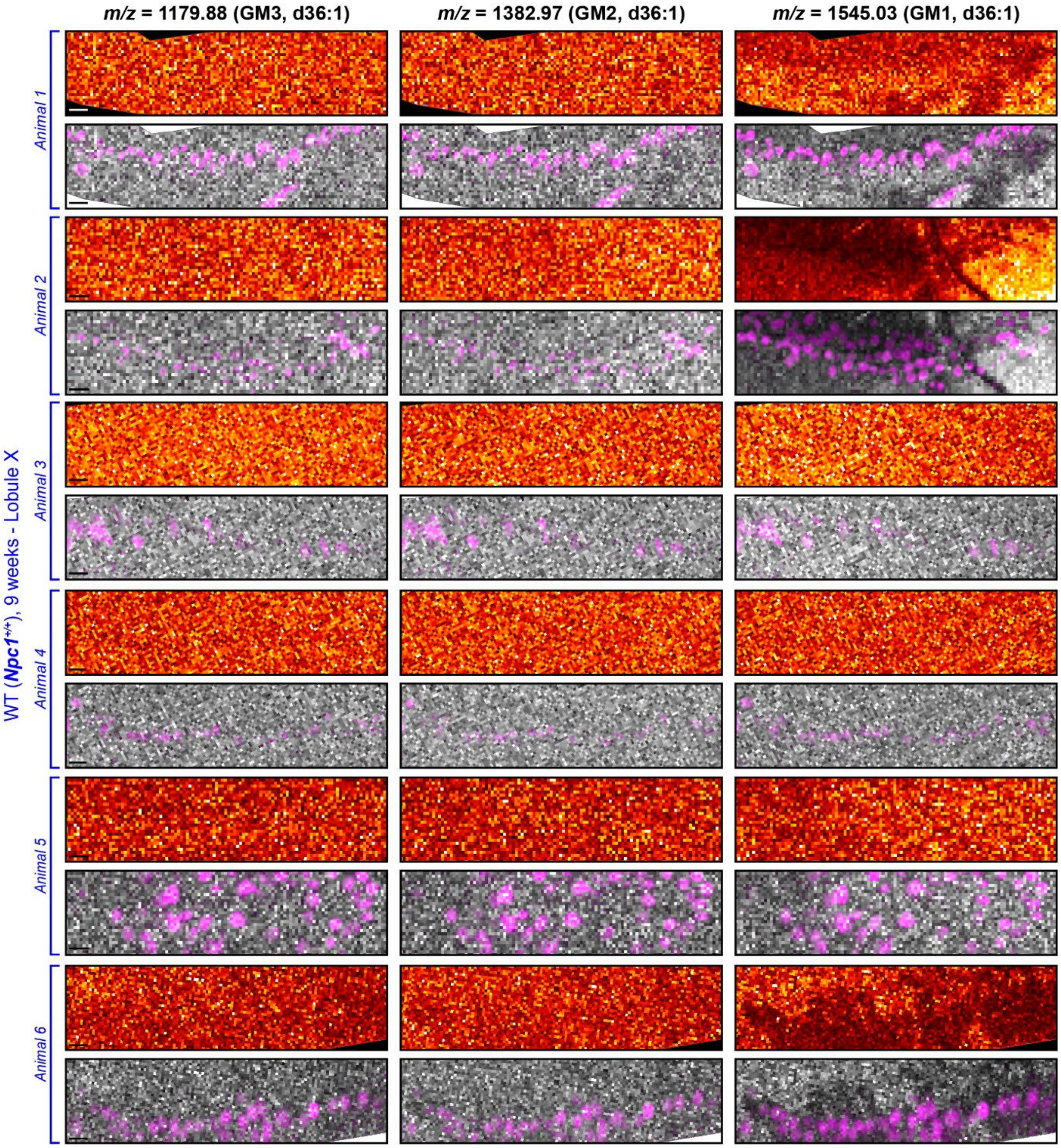

m/z = 1179.88 (GM3, d36:1)
m/z = 1382.97 (GM2, d36:1)
m/z = 1545.03 (GM1, d36:1)
WT (Npc1+/+), 9 weeks - Lobule X
Animal 1
Animal 2
Animal 3
Animal 4
Animal 5
Animal 6

**Supplementary Fig. S6: Spatial distribution of selected lipids in lobule X in WT (*Npc1*$^{+/+}$) mouse cerebellum.** iGAMSI protocol was performed on lobule X of 9-week WT mouse cerebellum for n=6 animals. For each animal: top row illustrates the spatial distribution of GM3 (*m/z* = 1179.88), GM2 (*m/z* = 1382.97), and GM1 (*m/z* = 1545.03), respectively. Bottom row illustrates the registered image overlays of IF images (pre-MSI) stained for calbindin (magenta) and MS images (grey). Scale bars, 25 µm (100 µm). Mass spectrometry images were collected on a rapifleX MALDI Tissuetyper (Bruker) with an instrument pixel size of 15 µm.

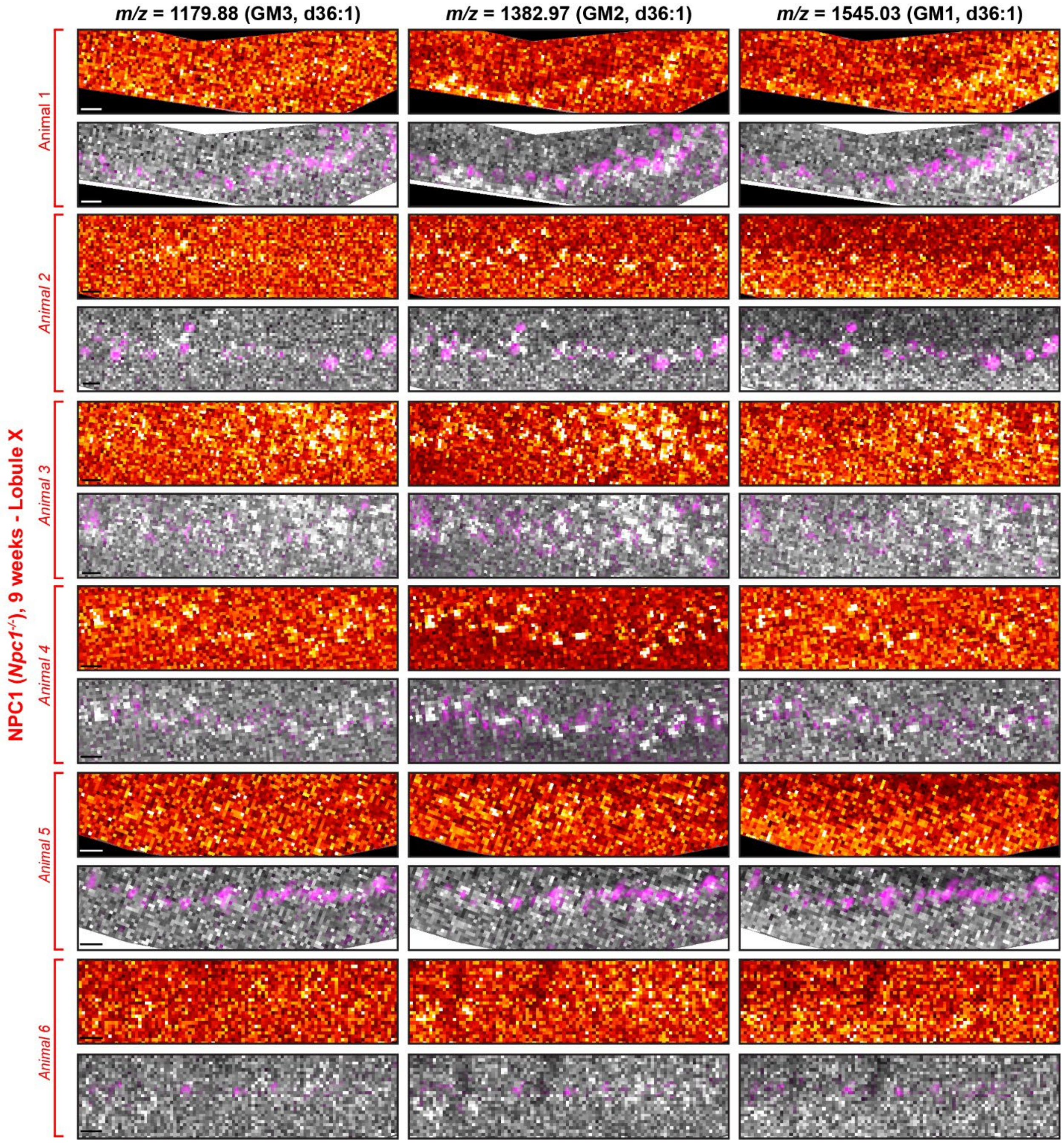
m/z = 1179.88 (GM3, d36:1)
m/z = 1382.97 (GM2, d36:1)
m/z = 1545.03 (GM1, d36:1)
NPC1 (Npc1-/-), 9 weeks - Lobule X
Animal 1
Animal 2
Animal 3
Animal 4
Animal 5
Animal 6

**Supplementary Fig. S7: Spatial distribution of selected lipids in lobule X in NPC1 (*Npc1*$^{-/-}$) mouse cerebellum.** iGAMSI protocol was performed on lobule X of 9-week NPC1 mouse cerebellum for n=6 animals. For each animal: top row illustrates the spatial distribution of GM3 (*m/z* = 1179.88), GM2 (*m/z* = 1382.97), and GM1 (*m/z* = 1545.03), respectively. Bottom row illustrates the registered image overlays of IF images (pre-MSI) stained for calbindin (magenta) and MS images (grey). Scale bars, 25 µm (100 µm). Mass spectrometry images were collected on a rapifleX MALDI Tissuetyper (Bruker) with an instrument pixel size of 15 µm.

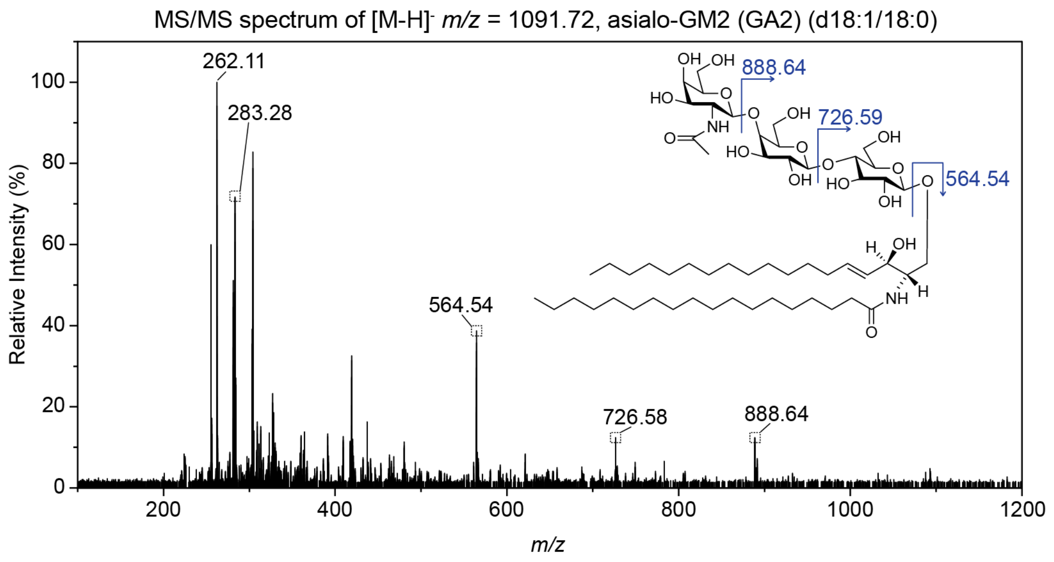


**Supplementary Fig. S8: Tandem mass spectrometry (MS/MS) validation of asialo-GM2 (GA2).** On-tissue MS/MS spectra of representative peak at *m/z* = 1091.72 from iGAMSI-processed mouse cerebellum. Spectra were collected using a timsTOF fleX QTOF Analyzer (Bruker).

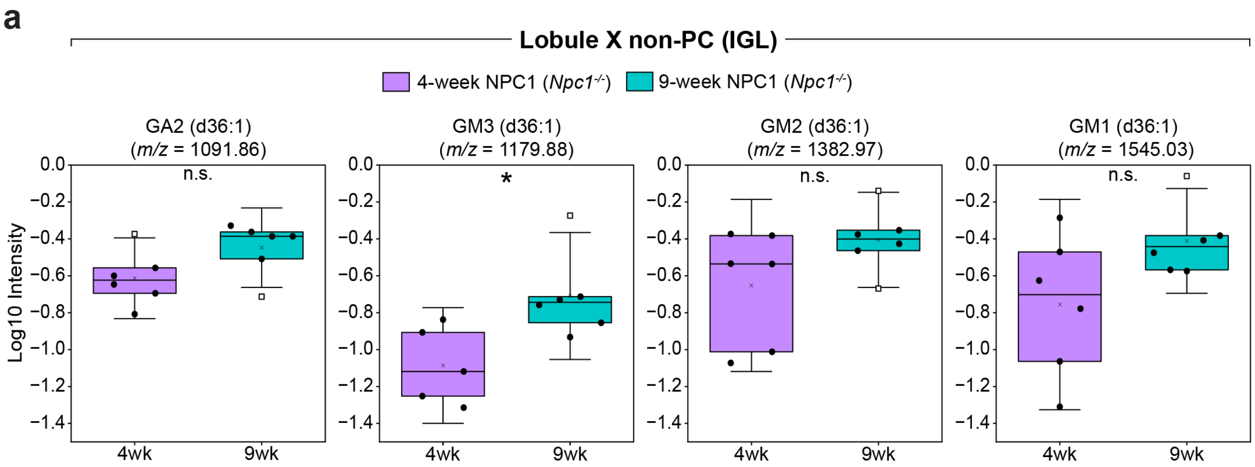


**Supplementary Fig. S9: Disease-progression- and lobule-dependent accumulation of sphingolipids within non-PCs in NPC1 cerebellum. a,** (Left to right) Relative abundance of GA2 d36:1 (*m/z* = 1091.86), GM3 d36:1 (*m/z* = 1179.88), GM2 d36:1 (*m/z* = 1382.97), and GM1 d36:1 (*m/z* = 1545.03) in non-PC (IGL) of lobule X in 4-week (purple) and 9-week (teal) NPC1 (*Npc*$^{-/-}$) mouse cerebella. Unpaired t-tests were performed (with Welch correction) with $p$ values of $p$ = 0.0751 (GA2), 0.0202 (GM3), 0.128 (GM2), and 0.085 (GM1). We note that for the 4-week GM3 plot in a and b, only five data points are shown as the log transform of one data point was an undefinable value.

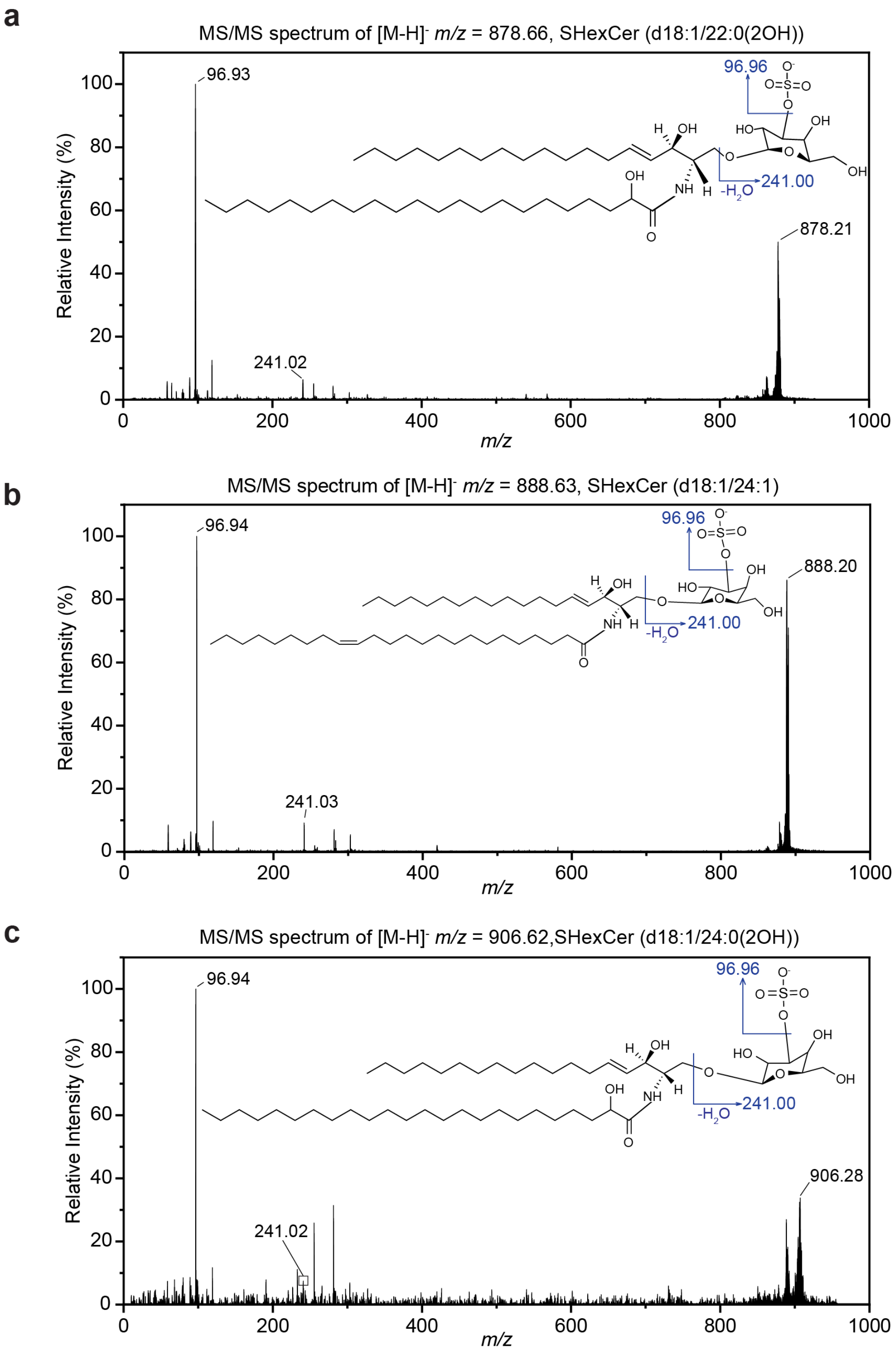


**Supplementary Fig. S10: Tandem mass spectrometry (MS/MS) validation of selected sulfatides.** On-tissue MS/MS spectra of representative peaks at *m/z* = **(a)** 878.66, **(b)** 888.63, **(c)** 906.62 from iGAMSI-processed mouse cerebellum. Spectra were collected using an AB SCIEX 4800 with instrument parameters set at CID gas off and MS/MS 2kV. SHexCer: sulfatide

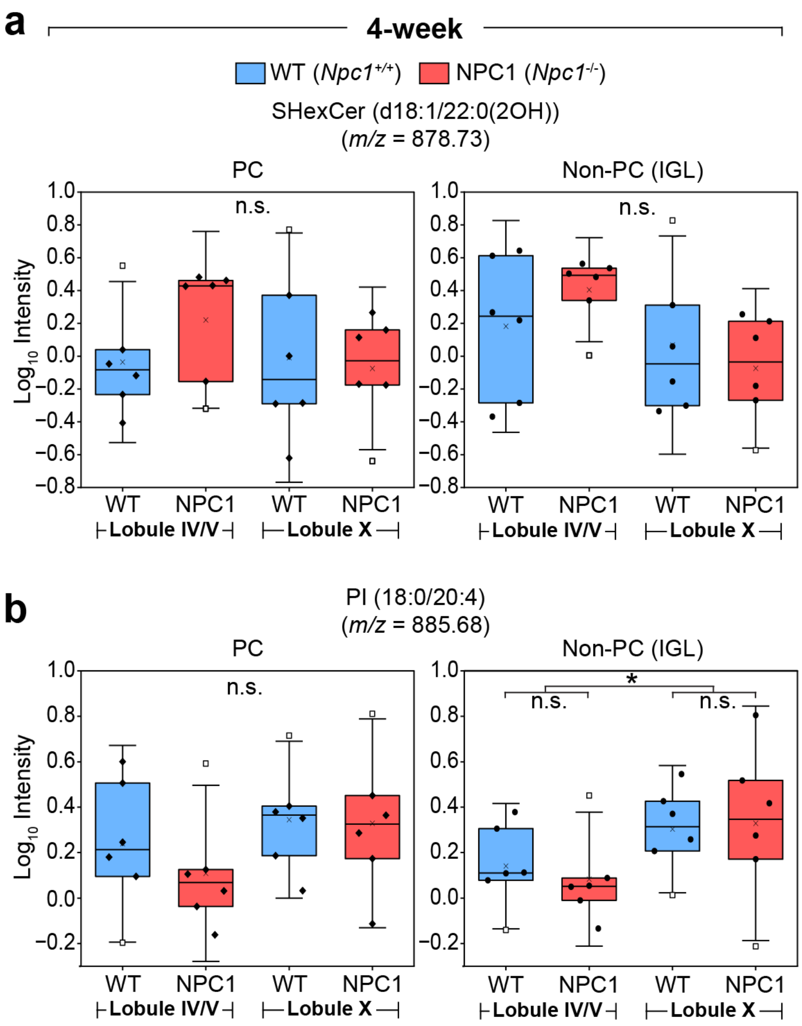


**Supplementary Fig. S11: Evaluation of lipidomic profiles of selected lipids across lobules and genotypes.** Box plots representing PCs (left, solid black diamonds) and non-PCs (right, solid black circles) of lobules IV/V and X in 4-week WT (blue) and NPC1 (red) mouse cerebellum for **(a)** SHexCer (d18:1/22:0(2OH)), (*m/z* = 878.73) and **(b)** PI (18:0/20:4), (*m/z* = 885.68). A 3-way ANOVA with Tukey's test was performed with a *p* value of $p = 0.048$ (non-PCs). For all plots, the y axis of all plots is the Log 10 ratio of the intensity of selected lipids to PE (18:0/22:6) (*m/z* = 790.66). Data are represented as box plots, where the ends of the whiskers represent standard deviation +/- 1.5 times the interquartile range, the upper line of the box represents the 75$^{th}$ percentile, the middle line represents the 50$^{th}$ percentile (median), the lower line represents the 25$^{th}$ percentile, the X represents the mean, and the open squares represent the individual values of the outlying data points. *$p < 0.05$, **$p < 0.01$, ***$p < 0.001$, ****$p < 0.0001$; n.s., not significant ($p \geq 0.05$).

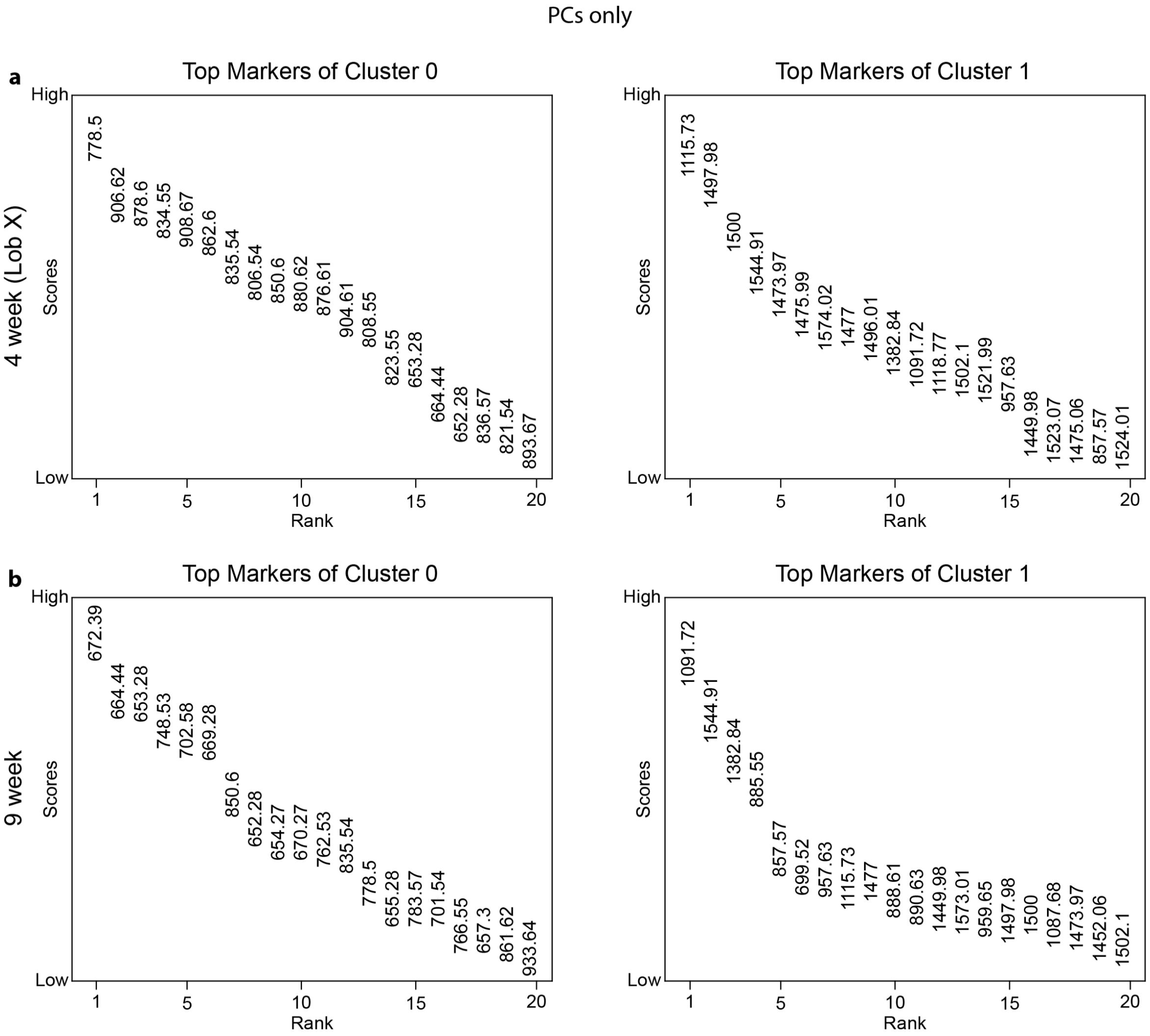


**Supplementary Fig. S12: Marker analysis of PCs. a,** Marker analysis of PCs lobule X (9-week) cerebella. **b,** Marker analysis of PCs lobule X (9-week) cerebella. Lipid features are arranged in descending order by their Wilcoxon scores (of which the values are indicated by the center of the *m/z* labels), with higher scores indicating stronger differential expression in the indicated cluster. The cluster indices (0, 1, …, n) correspond to the first, second, …, and (n+1)th clusters generated by Leiden clustering.

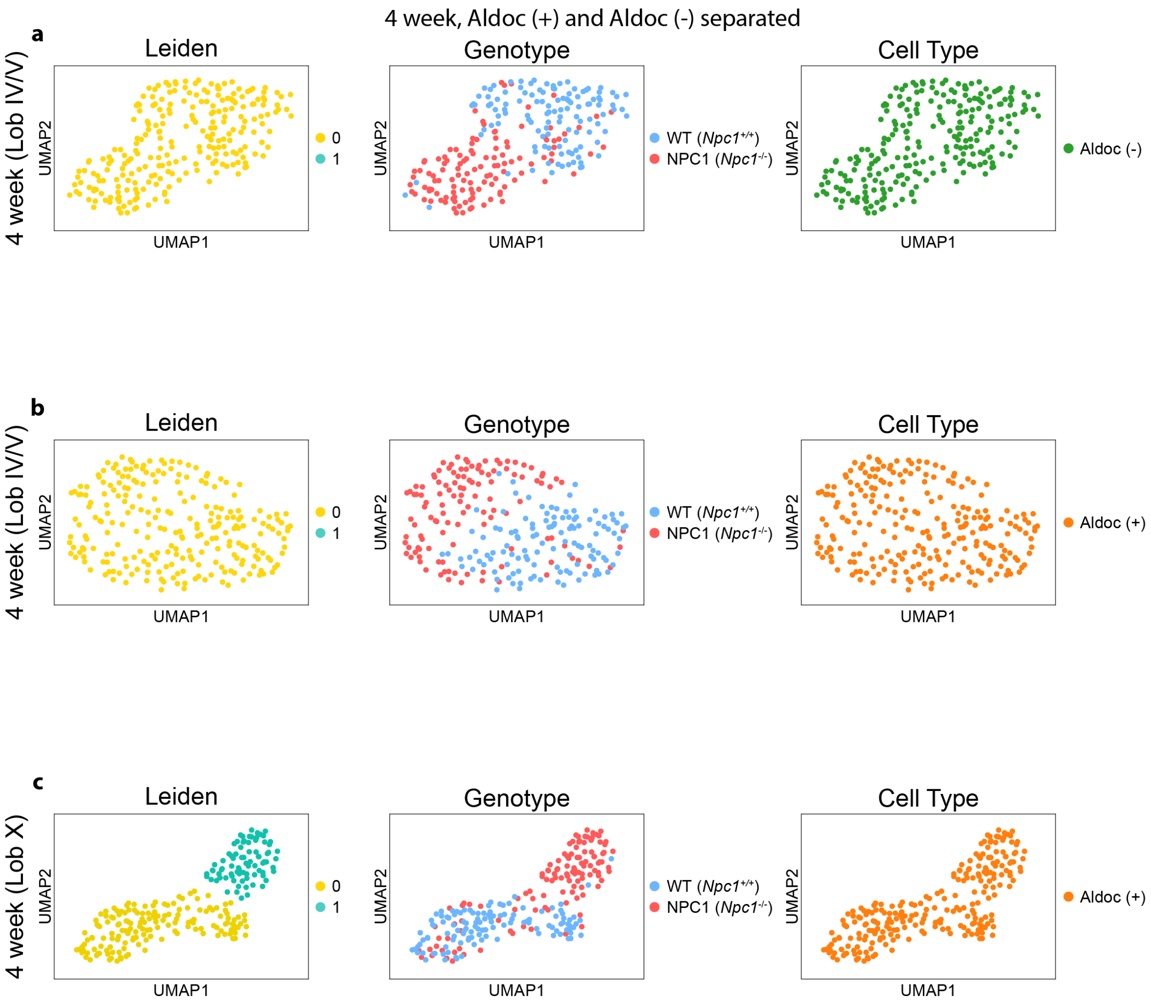


**Supplementary Fig. S13: Single-cell lipidomic clustering of PC subtypes of 4-week-old mice. a-c,** UMAP embedding of **(a)** aldolase-C-negative PCs of lobule IV/V (4-week), **(b)** aldolase-C-positive PCs of lobule IV/V (4-week), and **(c)** aldolase-C-positive PCs of lobule X (4-week) cerebella, annotated by (left to right) Leiden clusters, genotypes, and cell types. All UMAP embeddings were computed from neighbor graphs constructed from the data, with the Leiden clustering performed at a resolution of 1.0. AldoC (+): aldolase-C-positive; AldoC (-): Aldolase-C-negative. The cluster indices (0, 1, …, n) correspond to the first, second, …, and (n+1)th clusters generated by Leiden clustering.

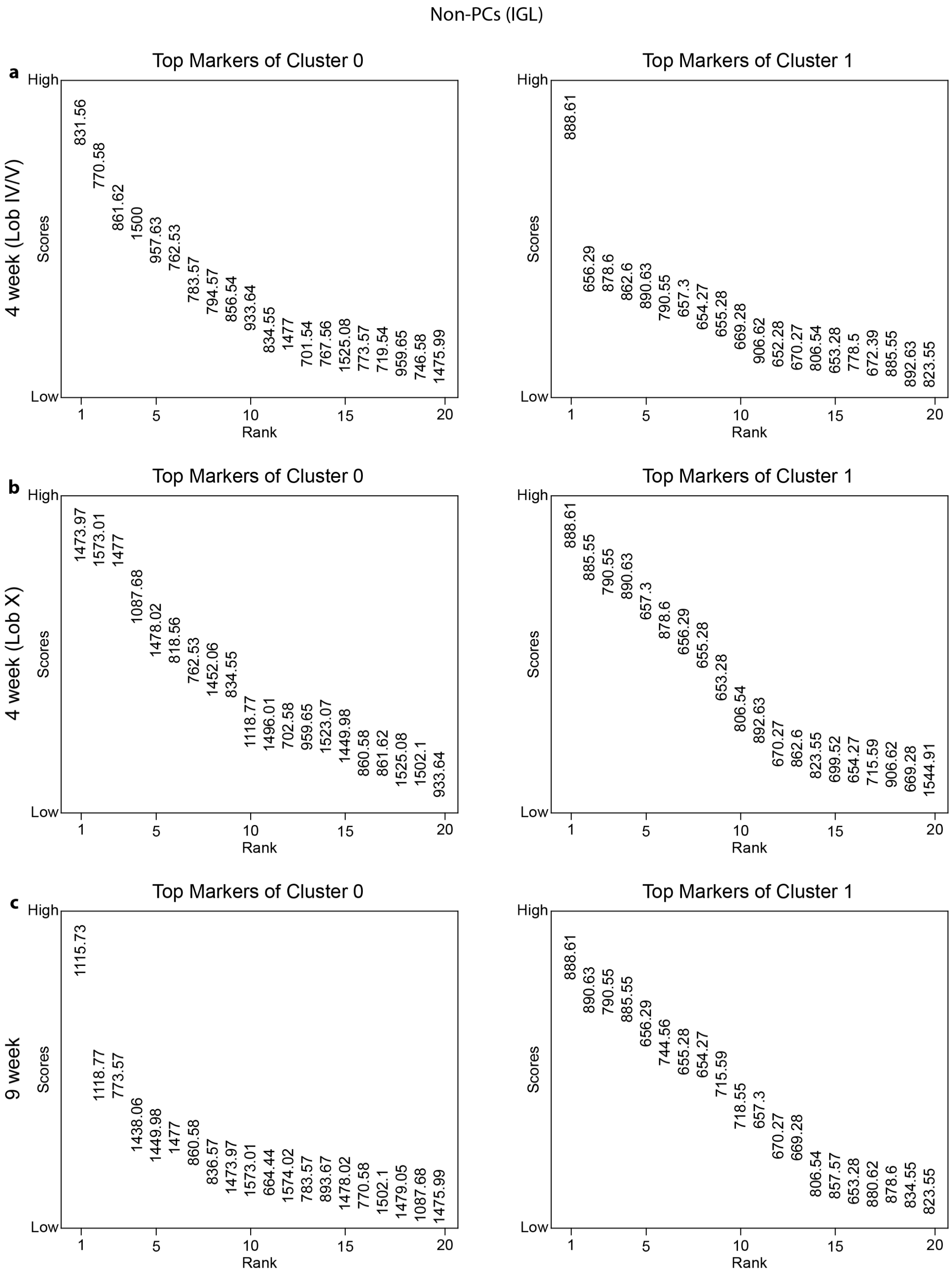
Non-PCs (IGL)
a
4 week (Lob IV/V)
Top Markers of Cluster 0
High
Low
Scores
Rank
831.56
770.58
861.62
1500
957.63
762.53
783.57
794.57
856.54
933.64
834.55
1477
701.54
767.56
1525.08
773.57
719.54
959.65
746.58
1475.99
Top Markers of Cluster 1
888.61
656.29
878.6
862.6
890.63
790.55
657.3
654.27
655.28
669.28
906.62
652.28
670.27
806.54
653.28
778.5
672.39
885.55
892.63
823.55
b
4 week (Lob X)
Top Markers of Cluster 0
1473.97
1573.01
1477
1087.68
1478.02
818.56
762.53
1452.06
834.55
1118.77
1496.01
702.58
959.65
1523.07
1449.98
860.58
861.62
1525.08
1502.1
933.64
Top Markers of Cluster 1
888.61
885.55
790.55
890.63
657.3
878.6
656.29
655.28
653.28
806.54
892.63
670.27
862.6
823.55
699.52
654.27
715.59
906.62
669.28
1544.91
c
9 week
Top Markers of Cluster 0
1115.73
1118.77
773.57
1438.06
1449.98
1477
860.58
836.57
1473.97
1573.01
664.44
1574.02
783.57
893.67
1478.02
770.58
1502.1
1479.05
1087.68
1475.99
Top Markers of Cluster 1
888.61
890.63
790.55
885.55
656.29
744.56
655.28
654.27
715.59
718.55
657.3
670.27
669.28
806.54
857.57
653.28
880.62
878.6
834.55
823.55

**Supplementary Fig. S14: Marker analysis of non-PCs (IGL). a,** Marker analysis of non-PCs (IGL) of lobule IV/V (4-week) cerebella. **b,** Marker analysis of non-PCs (IGL) of lobule X (4-week) cerebella. **c,** Marker analysis of PCs of lobule X (9-week) cerebella. Lipid features are arranged in descending order by their Wilcoxon scores (of which the values are indicated by the center of the *m/z* labels), with higher scores indicating stronger differential expression in the indicated cluster. The cluster indices (0, 1, …, n) correspond to the first, second, …, and (n+1)th clusters generated by Leiden clustering.

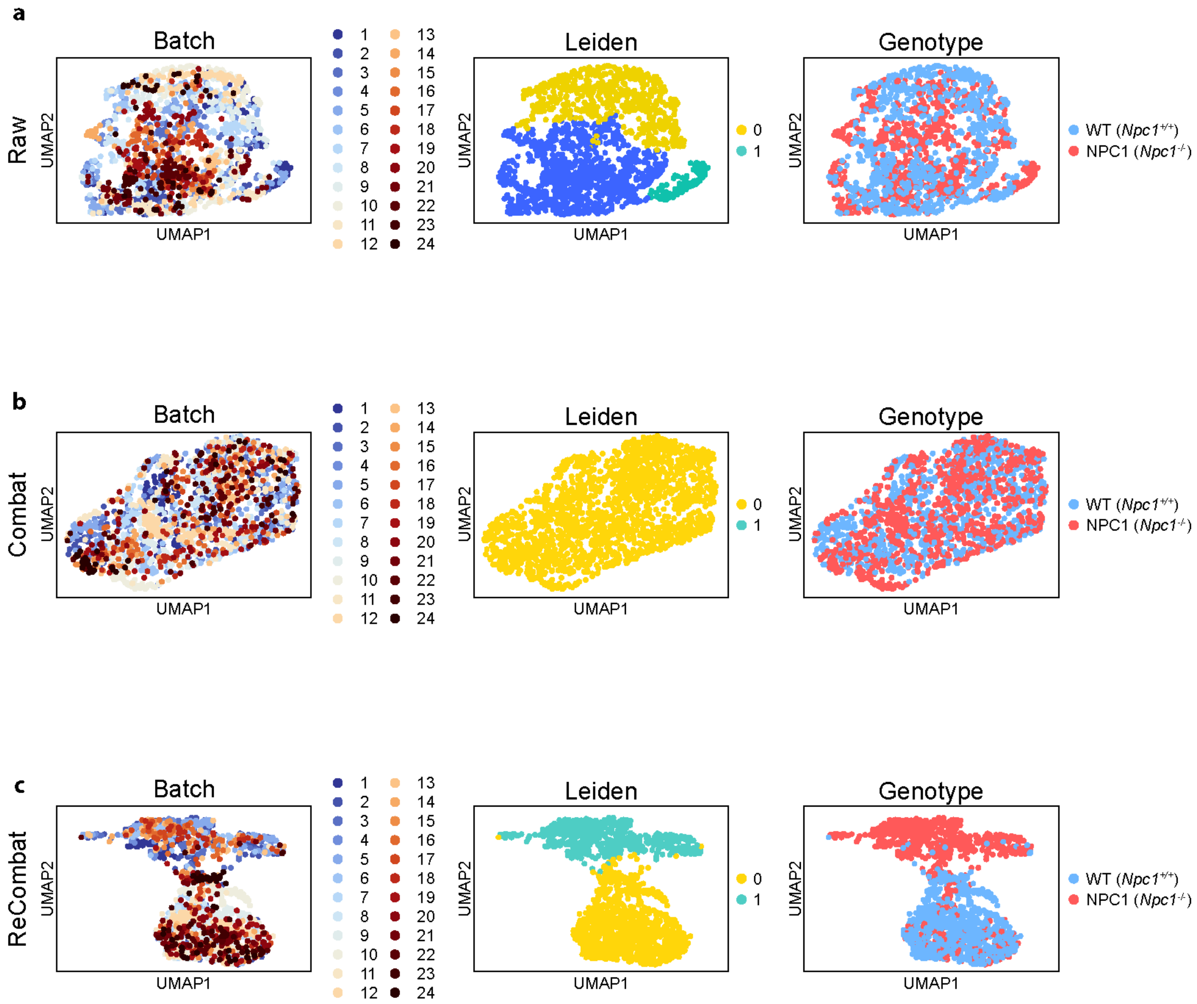


**Supplementary Fig. S15: Clustering analysis of data processed from different batch correction algorithms. a-c,** UMAP embedding of **(a)** raw data, **(b)** data processed from Combat, and **(c)** data processed from ReCombat, annotated by (left to right) batches, Leiden clusters, and genotypes. All UMAP embeddings were computed from neighbor graphs constructed from the data, with the Leiden clustering performed at a resolution of 1.0. The cluster indices (0, 1, …, n) correspond to the first, second, …, and (n+1)th clusters generated by Leiden clustering.

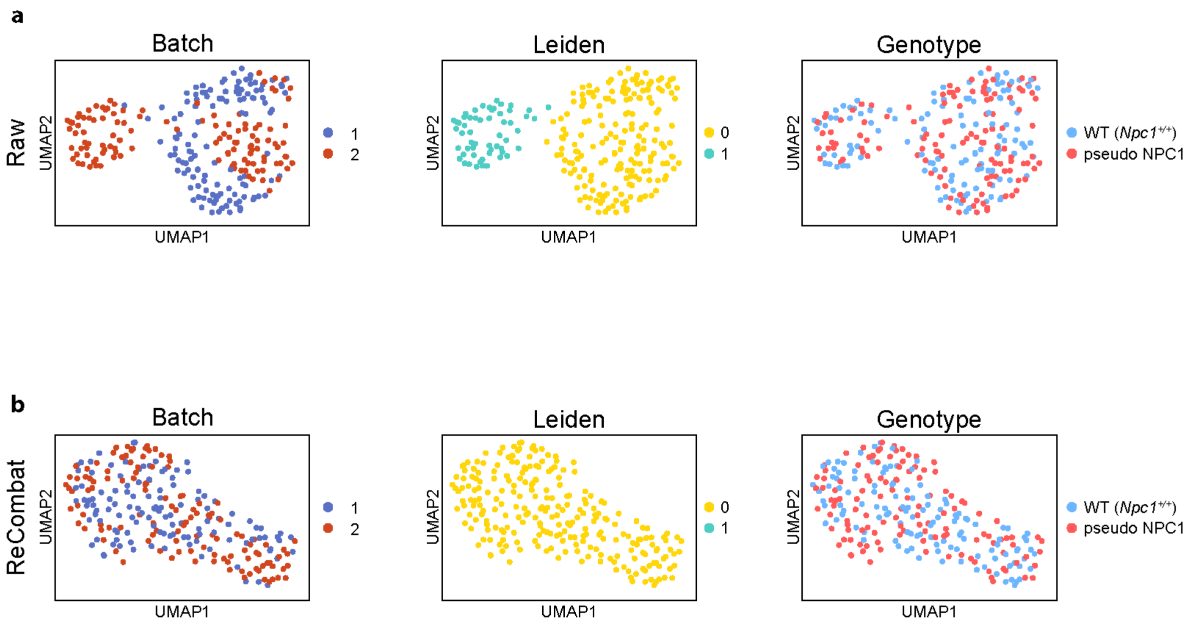


**Supplementary Fig. S16: Batch correction validation using pseudo-label. a-b,** UMAP embedding of **(a)** raw data with pseudo-label, and **(b)** data with pseudo-label processed from ReCombat, annotated by (left to right) batches, Leiden clusters, and genotypes. All UMAP embeddings were computed from neighbor graphs constructed from the data, with the Leiden clustering performed at a resolution of 1.0. The cluster indices (0, 1, …, n) correspond to the first, second, …, and (n+1)th clusters generated by Leiden clustering.

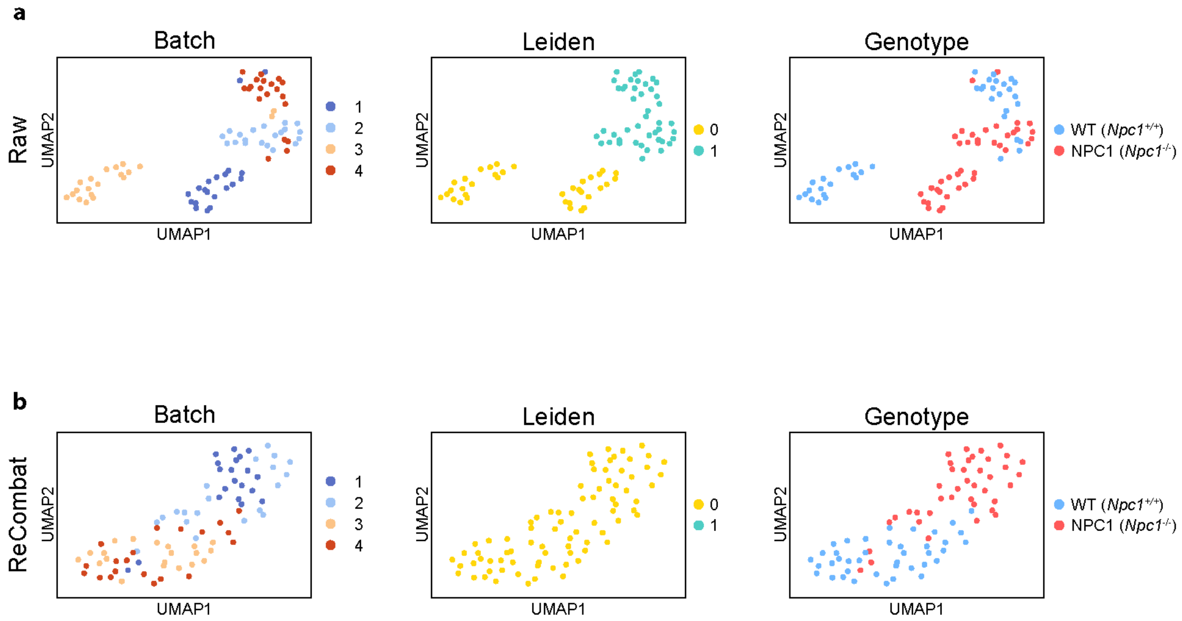


**Supplementary Fig. S17: Batch correction validation using data of technical replicates. a-b,** UMAP embedding of **(a)** raw data, and **(b)** data processed from ReCombat, annotated by (left to right) batches, Leiden clusters, and genotypes. All UMAP embeddings were computed from neighbor graphs constructed from the data, with the Leiden clustering performed at a resolution of 1.0. The cluster indices (0, 1, …, n) correspond to the first, second, …, and (n+1)th clusters generated by Leiden clustering.

#### Supplementary Tables

**Table S1: Primary antibodies used in this study.**

| Target | Host | Vendor | Cat. number |
|---|---|---|---|
| Calbindin | Guinea pig | Synaptic Systems | 214 004 |
| Zebrin-II (Monoclonal) | Rabbit | MilliporeSigma | ZRB1354-25UL |
| Zebrin-II (Polyclonal) | Rabbit | Thermo | PA5-27659 |
| TOMM20 | Rabbit | Abcam | ab186735 |
| NeuN | Chicken | Abcam | ab134014 |
| GFAP | Chicken | Abcam | Ab4674 |

**Table S2. Secondary antibodies used in this study.**

| Target | Host | Conjugated dye | Vendor | Cat. number |
|---|---|---|---|---|
| Guinea pig | Goat | Alexa Fluor 568 | ThermoFisher | A11075 |
| Rabbit | Goat | Alexa Fluor 488 | ThermoFisher | A11008 |
| Rabbit | Goat | ATTO 647N | Rockland | 611-156-122 |
| Chicken | Goat | Alexa Fluor 488 | ThermoFisher | A11039 |
| Chicken | Goat | Alexa Fluor 568 | ThermoFisher | A11041 |

**Table S3. Optimization of homogenization conditions for iGAMSI.**

| Conditions | Protease | Time | Temperature | Lipid retention (based on optical scattering of the white matter region) | Sample-hydrogel composite integrity |
|---|---|---|---|---|---|
| 1 | Trypsin | 2 days | Room temperature (RT; ~20-22°C) | High | Tissue tearing and wrinkling upon expansion |
| 2 | Proteinase K | Overnight | RT | Low | Intact and expanded well |
| 3 | Proteinase K | 6 hours | RT | Middle | Intact and expanded well |
| 4 | Proteinase K | 4 hours | RT | High | Intact and expanded well |

| | | | | | |
|---|---|---|---|---|---|
| 5 | Proteinase K | 2 hours | RT | High | Tissue tearing and wrinkling upon expansion |

**Table S4.** ***p*** **values for 2-way ANOVA.**

| Peak value (*m/z*) | Genotype level | Aldolase C expression level |
|---|---|---|
| **1091.86 (GA2)** | 0.0247 | 0.593 |
| **1179.88 (GM3)** | 0.254 | 0.684 |
| **1382.97 (GM2)** | 0.0003 | 0.718 |
| **1545.03 (GM1)** | 0.380 | 0.844 |

**Table S5.** ***p*** **values for 2-way ANOVA (PCs)**

| Peak value (*m/z*) | Genotype level | Lobule level |
|---|---|---|
| 888.74 SHexCer, (d18:1/24:1) | 0.907 | 0.156 |
| 906.74 SHexCer (d18:1/24:0(2OH)) | 0.852 | 0.285 |
| 878.73 (d18:1/22:0(2OH)) | 0.553 | 0.407 |
| 885.68, PI (18:0/20:4) | 0.519 | 0.158 |

**Table S6.** ***p*** **values for 2-way ANOVA (non-PCs)**

| Peak value (*m/z*) | Genotype level | Lobule level |
|---|---|---|
| 888.74 SHexCer, (d18:1/24:1) | 0.527 | 0.012 |
| 906.74, SHexCer (d18:1/24:0(2OH)) | 0.279 | 0.030 |
| 878.73, SHexCer (d18:1/22:0(2OH)) | 0.788 | 0.060 |
| 885.68, PI (18:0/20:4) | 0.870 | 0.0478 |
